# Sleep promotes cortical neuronal autophagy for intracellular clearance

**DOI:** 10.64898/2026.11.08.764098

**Authors:** Lin Yang, Xue Zhang, Jiaqi Li, Xueying Liu, Danqian Liu

**Affiliations:** Institute of Neuroscience, Center for Excellence in Brain Science and Intelligence Technology, Chinese Academy of Sciences, Shanghai 200031, China; University of Chinese Academy of Sciences, Beijing 100049, China; Sino-Danish College at University of Chinese Academy of Sciences, Beijing 100049, China

**Keywords:** sleep, neuronal autophagy, sleep function, neurodegeneration

## Abstract

Sleep is essential for brain health, yet how it maintains neuronal homeostasis at the cellular level remains poorly understood. Here, we show that extended wakefulness selectively suppressed neuronal autophagic flux in the cortex of mice, whereas sleep rapidly restored it, accompanied by a shift in autophagic cargo from mitochondrial components to synaptic proteins. Sleep upregulated ATG7 expression in the cortex, involving both lysosomal degradation and a putative noncanonical secretory route. Moreover, low neuronal activity is associated with efficient flux, as Gq-mediated neuronal activation impaired autophagy and promoted intracellular amyloid-β accumulation. Notably, closed-loop optogenetic stimulation of lysosomal acidification, exclusively during sleep, reduced amyloid pathology in 5xFAD Alzheimer’s mice, whereas wake-targeted stimulation worsened pathology. Together, these findings demonstrate that sleep serves as a physiological gatekeeper for cortical neuronal autophagy, and highlight the importance of brain-state timing for enhancing intracellular proteostasis in neurodegenerative disease.

## Main

Sleep is an innate, recurring behavioral state indispensable for brain health. Acute sleep deprivation impairs learning and memory, and chronic sleep deprivation compromises neuronal integrity and contributes to psychiatric and neurodegenerative disorders, and in extreme cases, can be fatal [1–7]. Sleep has been proposed to support a broad range of restorative processes, from synaptic and metabolic homeostasis to molecular repair and waste clearance [2, 8, 9], yet the cellular mechanisms through which sleep supports these functions remain largely unknown. Among these, intracellular clearance is particularly important for long-lived neurons, which rely on continuous removal of damaged proteins and organelles to maintain proteostasis [2, 10]. Sleep-dependent dynamics of glymphatic and cerebrospinal fluid was reported to facilitate extracellular waste removal [11–13]. By contrast, how sleep regulates intracellular clearance remains much less understood.

Autophagy, a multi-stage cellular pathway spanning initiation, cargo selection, autophagosome formation, and lysosomal degradation, is central to neuronal homeostasis [14–17]. While extensively characterized under stress and disease conditions [18], autophagy is also regulated by physiological states such as fasting and circadian rhythms [19, 20]. Sleep-wake transitions occur rapidly and involve heterogeneous changes across the brain [21, 22], making this dynamic process difficult to assess using static assays [23–25]. Recent studies in Drosophila have shown that neuronal autophagy varies across the circadian cycle, and is further influenced by prior wake history [26, 27]. In rodents, however, sleep deprivation studies have reported inconsistent changes in static autophagy markers, interpreted as either increased or decreased autophagy [28]. Resolving these ambiguities therefore requires systematic *in vivo* measurements of autophagic flux with cell-type resolution [29, 30].

In this study, we combined cell-type-resolved autophagic flux reporters, electron microscopy, proteomic profiling of LC3-immunopurified autophagic vacuoles, and genetic manipulation, to assess brain autophagy dynamics across sleep-wake states. We found that sleep selectively promotes autophagic flux in cortical excitatory neurons, through both molecular regulation of the autophagic machinery and reduced neuronal activity. Furthermore, using optogenetic control of lysosomal acidification, we provide causal evidence that boosting lysosomal function is beneficial specifically during sleep, when endogenous autophagic flux is high. Thus, these results establish sleep-autophagy alignment as critical for effective intracellular clearance and highlight brain-state timing as an important consideration for enhancing proteostasis in neurodegenerative disease.

### Sleep selectively promotes autophagic flux in cortical excitatory neurons

To monitor brain autophagic flux, we used transgenic mice expressing the tandem fluorescent reporter CAG-RFP-EGFP-LC3 (CAG-tfLC3) [30, 31], and combined it with immunostaining (e.g., pan-neuronal marker NeuN), to identify distinct cell populations (**Figure 1A,B**). Mice were subjected to 6 h of sleep deprivation (SD; ZT0– 6), SD followed by 2 h of recovery sleep (SR), or undisturbed sleep (Ctrl; **Figure 1A; Supplementary Figure 1A,B**). We first focused on the cortex to examine state regulation of autophagic flux. Within neuronal somata, neutral LC3^+^ autophagic vacuoles (AVs) appear yellow (RFP⁺GFP⁺), whereas acidified AVs quench GFP fluorescence and appear red (RFP⁺GFP⁻; **Figure 1B**) [30, 31]. In natural sleeping mice at ZT 6 or ZT 8 (Ctrl), NeuN⁺ cortical neurons displayed abundant somatic tfLC3 puncta (30.51 ± 2.72 puncta/neuron) and a high acidification ratio (∼0.8, red/total puncta ratio; **Figure 1B–E**). SD markedly suppressed both measures, reducing tfLC3 puncta to 5.78±0.53 puncta/neuron and acidification ratio to ∼0.3 (P<0.0001, one-way ANOVA; **Figure 1B–E**), accompanied by increased p62 accumulation (**Supplementary Figure 1C**). Notably, 2 h of SR robustly reversed these changes, restoring both puncta abundance and acidification to control levels (P<0.0001; **Figure 1B–E**). Similar increases in AV abundance were observed in proximal apical dendrites (**Supplementary Figure 1D**). A time-course comparison at 0.5, 1, and 2 h of SR further showed that 0.5 h of SR (∼15 mins of sleep), was sufficient to enhance autophagic flux, with AV acidification reaching a plateau while AV abundance continued to increase with longer sleep (**Supplementary Figure 1E**).

**Figure 1.**
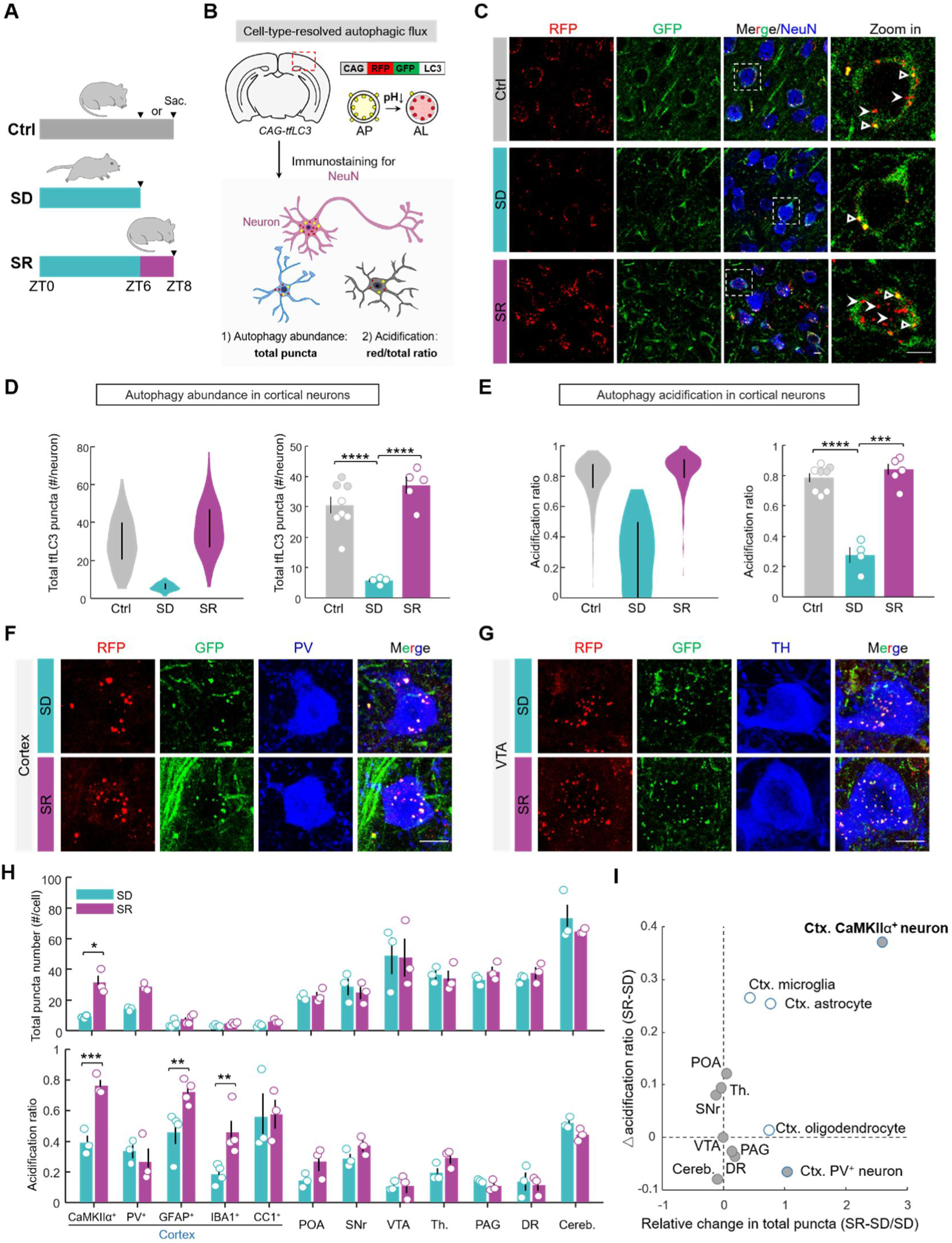
Sleep selectively promotes autophagy flux in cortical excitatory neurons. A. Experimental timeline. Mice were left undisturbed in their home cage (Ctrl), subjected to 6 h of sleep deprivation (SD), or allowed 2 h of sleep recovery (SR) after SD. ZT: zeitgeber time. Arrowheads indicate tissue collection. Sac., sacrifice. B. ***Top:*** Schematic of the ubiquitously expressed CAG-driven tandem-fluorescent LC3 (tfLC3) reporter transgene. ***Bottom:*** Detection of tfLC3 signals in cortical neurons by immunostaining for the pan-neuronal marker NeuN. Total tfLC3 puncta within each cell indicate overall autophagic vacuole abundance, and the red-to-total puncta ratio reflects AV acidification. C. Representative fluorescence images of tfLC3 signals and NeuN immunostaining in mouse cortex across ctrl, SD and SR conditions. Insets show zoom-in views of individual neurons. Solid arrowheads indicate representative acidified puncta (RFP⁺ only, red); open triangles indicate neutral puncta (RFP⁺&GFP⁺, yellow). D. ***Left:***Violin plot showing the distribution of tfLC3 puncta numbers across n= ∼130-270 visual cortical neurons pooled from multiple mice (N = 8, 4, and 5 mice for Ctrl, SD, and SR, respectively). Vertical lines indicate interquartile range (IQR). ***Right:***Mouse-level quantification. Each dot represents one mouse; in the Ctrl group, filled and open dots denote ZT6 (N = 3; matched to SD) and ZT8 (N = 5; matched to SR), respectively. E. Similar to (D), but showing the neuronal acidification ratio (red/total tfLC3 puncta). F. Representative images of tfLC3 signals in parvalbumin (PV)-expressing cortical interneurons, under SD and SR conditions. G. Similar to (F), but in TH-expressing dopaminergic neurons in the ventral tegmental area (VTA). H. Quantification of tfLC3 puncta abundance (**top**) and acidification ratio (**bottom**) across cell populations under SD and SR conditions (N=3 or 4 mice/group). Cortical cell populations were identified by CaMKII, PV, GFAP, IBA1, or CC1 immunostaining. Subcortical neuronal populations were identified by NeuN immunostaining, but TH immunostaining was used for VTA. Each dot represents data from one mouse. POA, preoptic area; SNr, substantia nigra pars reticulata; Th., thalamus; PAG, periaqueductal gray; DR, dorsal raphe; Cereb., cerebellum. I. Summary of SR-induced changes in puncta abundance and acidification. Each point represents one cell population. Blue outlines indicate cortical populations, and gray fill indicates neuronal populations. Note that the CaMKIIα-expressing cortical excitatory neurons showed the largest changes in both measures. Error bars represent SEM. One-way ANOVA with Bonferroni’s correction was used for mouse-level comparisons in (D), (E); two-way ANOVA with Bonferroni’s correction was used for comparisons within each cell population in (H). *P < 0.05; **P <0.01; ***P<0.001; ****P < 0.0001. Scale bars, 5 μm in (C); 10 μm in (F-G).

We next systematically compared tfLC3 signals across major cortical cell types and multiple subcortical neuronal populations (**Figure 1F–I; Supplementary Figure 1F– I**). Notably, CaMKIIα⁺ cortical excitatory neurons showed the largest increase in both AV abundance and acidification during SR, with acidified AV puncta increasing more than sevenfold, from 3.4 ± 0.5 to 24.2 ± 4.8 puncta per cell (mean ± SEM; **Figure 1H,I; Supplementary Figure 1I**). Parvalbumin (PV)⁺ inhibitory neurons showed a tendency toward increased AV abundance during SR, mainly due to increased neutral AVs (9.7 vs 21.5 puncta/cell), suggesting limited acidification (**Figure 1F,H,I; Supplementary Figure 1H,I**). Cortical glial cells, however, contained relatively few AVs (∼5 puncta/cell), with astrocytes and microglia showing increased AV acidification (**Figure 1H,I; Supplementary Figure 1F-I**). In contrast, subcortical neurons generally showed much higher AV abundance (>30 puncta/cell) than cortical populations and remained largely unchanged across sleep-wake states, although cerebellar Purkinje cells showed reduced acidified AVs during SR (**Figure 1G–I; Supplementary Figure 1H,I**). Thus, sleep-wake modulation of autophagy was highly cell-type specific, with cortical excitatory neurons showing the most prominent increase during sleep.

Because puncta-based tfLC3 readouts may be confounded by punctate autofluorescence (e.g., lipofuscin) [32], we used orthogonal approaches to assess cortical autophagic flux [25]. We generated a Cre-dependent AAV expressing a second reporter GFP-LC3-RFP, in which cytosolic RFP serves as an internal control and a lower GFP/RFP ratio indicates greater acidification [33], and injected it into *CaMKIIα^Cre^* mice to label cortical excitatory neurons (**Supplementary Figure 2A**). In SD neurons, GFP/RFP fluorescence ratio was significantly higher (P=0.02, one-way ANOVA), with neurons appearing predominantly yellow (**Supplementary Figure 2B,C**), consistent with impaired acidification (**Figure 1E**). SR shifted the fluorescence toward red, reflecting GFP quenching and partial restoration of acidification (**Supplementary Figure 2B,C**). Moreover, ultrastructural examination of cortical neurons by electron microscopy (**Supplementary Figure 2D-F**) showed that autolysosomes (ALs) were significantly more abundant under sleep conditions (Ctrl vs. SD: P=0.016; SR vs. SD: P=0.0065; **Supplementary Figure 2E**), while autophagosomes (APs) remained largely unchanged (**Supplementary Figure 2D**), confirming enhanced AV maturation.

Taken together, these findings show that sleep selectively enhances autophagic flux in cortical excitatory neurons, within minutes, with increases in AV abundance and acidification detectable within minutes of sleep.

### Sleep preferentially directs synaptic cargo into autophagic vacuoles

We next asked whether the protein composition of AV cargo changes across sleep-wake states. To address this, we isolated LC3⁺ AVs for proteomic analysis using a modified two-step purification strategy [34–36]. Forebrain homogenates from SD or SR were first subjected to density-gradient centrifugation to obtain AV-enriched fractions (**Figure 2A,B**), while omitting glycyl-L-phenylalanine 2-naphthylamide (GPN) to retain acidic ALs together with APs (**Supplementary Figure 3A**) [36, 37]. The obtained fractions were then purified by LC3 immunoprecipitation (IP) to remove LC3-negative contaminants [38], with parallel IgG IP serving as a negative control for nonspecific binding (N=3 biological replicates; **Figure 2B; Supplementary Figure 3B**). The purified AVs were subsequently treated with proteinase K to remove externally associated proteins [34], thereby enriching membrane-protected AV content for quantitative proteomics (**Figure 2A,B**).

**Figure 2.**
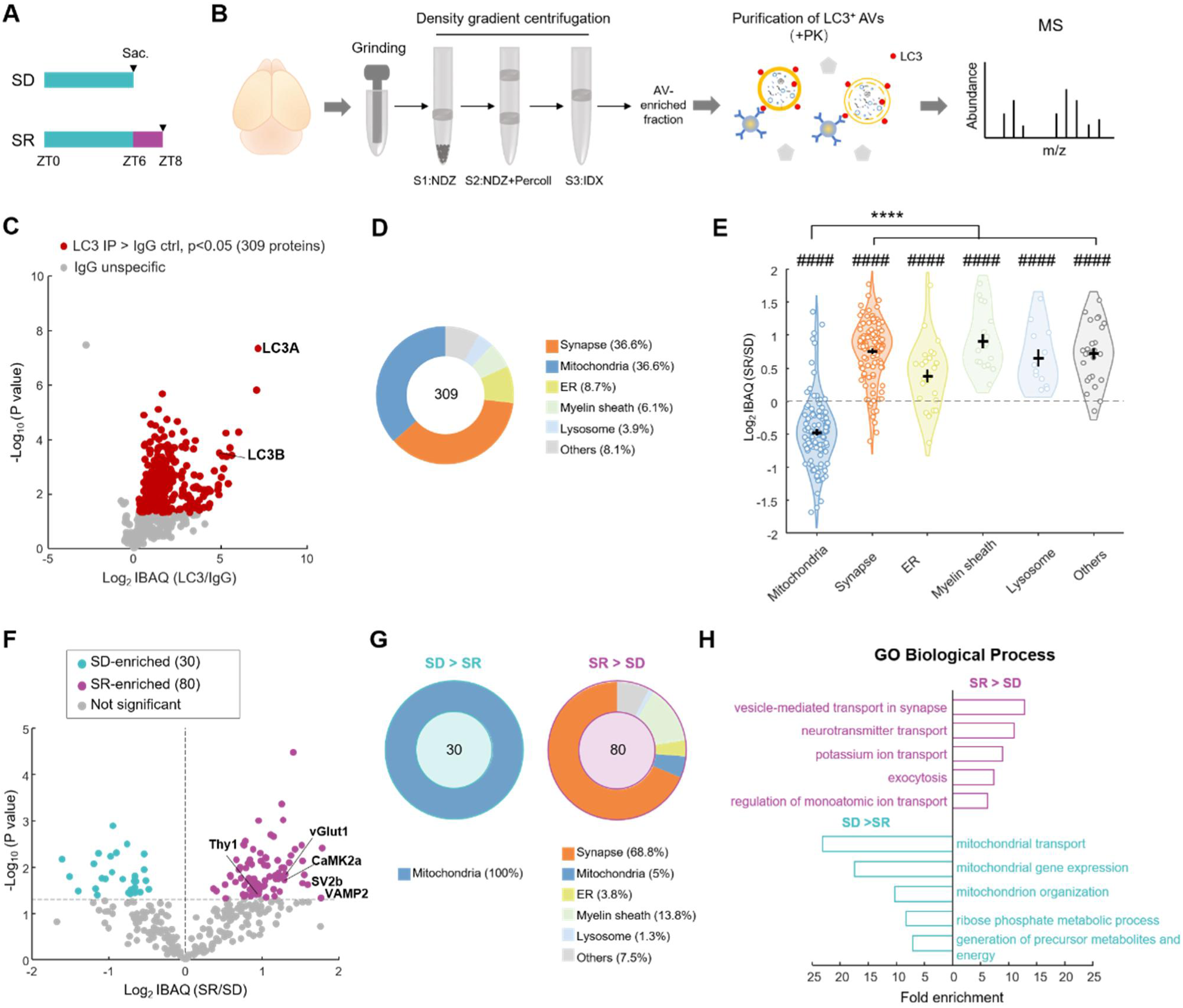
LC3⁺ autophagic vacuole proteomics reveals preferential targeting of synaptic cargo during sleep. A. Experimental timeline. B. Schematic of LC3⁺ AV purification and proteomics. Forebrain homogenates were subjected to sequential density-gradient centrifugation to obtain an AV-enriched fraction, followed by LC3 immunoprecipitation and proteinase K (PK) treatment to enrich membrane-protected AV cargo for quantitative mass spectrometry (MS). C. Volcano plot comparing proteins detected in LC3-IP and IgG controls. Proteins significantly enriched in LC3-IP are shown in red (P < 0.05, unpaired t test); LC3A and LC3B are indicated. D. Donut plot showing the subcellular distribution of the AV cargo proteome. E. Relative abundance of proteins across subcellular categories in SR versus SD, shown as log2 iBAQ(SR/SD). Each dot represents one protein (synapse, n = 113; mitochondria, n = 113; ER, n = 27; myelin sheath, n = 18; lysosome, n = 12; others, n = 26). Error bars: SEM. ****P < 0.0001, Welch’s one-way ANOVA followed by Games–Howell correction for across-category comparisons; ####P < 0.0001, one-sample t-test versus 0 for each category. F. Differential analysis of AV cargo between SR and SD. SR-enriched proteins are shown in magenta and SD-enriched proteins in cyan (P < 0.05, unpaired t-test); nonsignificant proteins are shown in gray. Selected synaptic proteins are labeled. G. Subcellular composition of SD- and SR-enriched AV proteins. H. Gene Ontology (GO) biological-process enrichment analysis of SR- and SD-enriched AV proteins.

Across groups, we identified 608 proteins by LC-MS analysis (**Data S1**). Of these, 309 proteins were significantly enriched in the LC3-IP compared with IgG controls (P < 0.05; **Figure 2C; Data S1**), and were defined as the AV cargo proteome. LC3A and LC3B were among the most strongly enriched proteins (>5-fold over IgG), supporting the specificity of the purification (**Figure 2C**). This AV cargo proteome mapped broadly to synaptic, mitochondrial, ER, myelin, and lysosomal compartments (**Figure 2D**). Among these, synaptic and mitochondrial proteins were the most abundant, each accounting for ∼37% of the AV proteome (**Figure 2D**). Notably, mitochondrial proteins showed higher relative abundance in SD AVs, whereas synaptic proteins were more abundant in SR AVs (**Figure 2E**).

This state distinction was further supported by differential analysis, which showed more AV cargo proteins enriched in SR than in SD (80 vs. 30; P < 0.05; **Figure 2F; Supplementary Figure 3C**). Of the 30 SD-enriched proteins, all were mitochondrial, whereas 68.8% of the 80 SR-enriched proteins were synaptic and only five were mitochondrial (**Figure 2G; Supplementary Figure 3D-H**). Among the SR-enriched synaptic proteins, a larger fraction localized to presynaptic than postsynaptic compartments (30.9% vs. 14.5%; **Supplementary Figure 3G**), and Gene Ontology (GO) analysis highlighted synaptic vesicle and neurotransmitter transport (**Figure 2H**). Despite the use of whole-forebrain AV preparations, several proteins predominantly expressed in cortical excitatory neurons, including VGLUT1, CaMKIIα and Thy1 [39–41], were more abundant in SR AVs (**Figure 2F; Supplementary Figure 3I**). Consistently, glutamatergic components (P<0.0001, Fisher’s exact test), but not GABAergic components, were significantly overrepresented in SR-enriched AV proteins (**Supplementary Figure 3I,J**). This preferential targeting of excitatory synaptic components is in line with the cortical excitatory neuron bias observed above (**Figure 1**), and may be associated with the restoration of excitatory synaptic function during sleep [8, 42–46].

Collectively, sleep not only enhances autophagic flux in cortical excitatory neurons, but also shifts autophagic cargo toward synaptic proteins.

### ATG7 knockdown attenuates sleep-associated neuronal autophagy

To determine how sleep regulates cortical autophagy, we examined the expression of core autophagy, lysosomal, and endolysosomal trafficking genes by qPCR [25, 47] (**Figure 3A**). Among core components of the canonical autophagy machinery [48], *Atg7* and *Beclin1* were reduced after SD and restored during SR. Atg5, however, showed the opposite pattern and increased during SD (**Figure 3A,B**), suggesting differential regulation of autophagy genes [49–51]. While some vesicular trafficking genes, including *Cd63* and *Rab6b,* also increased during SR, most lysosomal genes remained relatively stable (**Figure 3A,B**). Among these changes, Atg7 showed the strongest sleep-dependent upregulation in the cortex, whereas comparable changes were not observed in the subcortical regions examined (**Figure 3B**).

**Figure 3.**
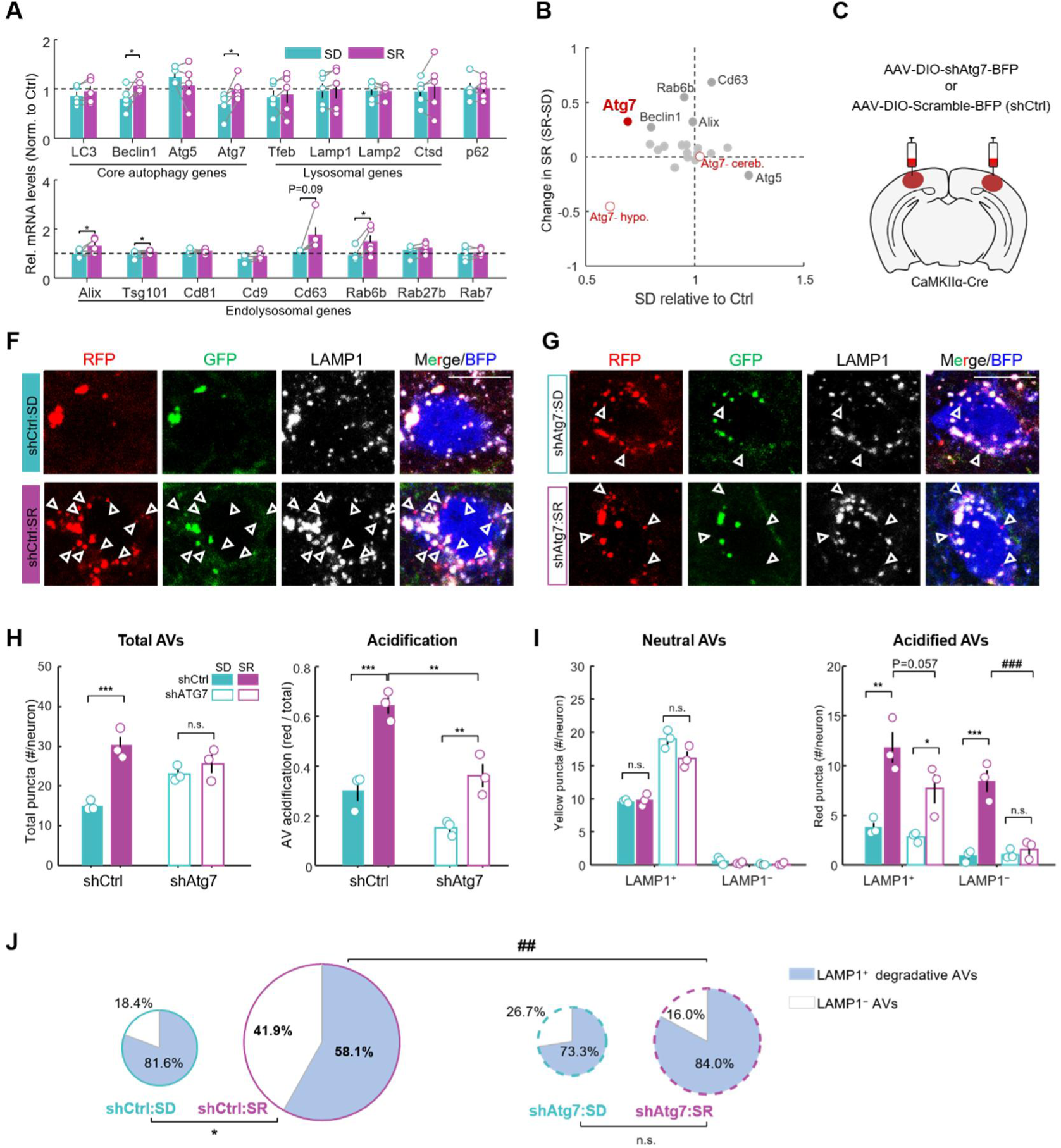
ATG7 is required for sleep-driven autophagy in cortical excitatory neurons. A. Cortical mRNA expression of core autophagy, lysosomal, and endolysosomal trafficking genes in SD and SR mice, normalized to Ctrl (N=5 mice/group). Dashed line indicates Ctrl level at 1. B. Summary of changes in gene expression during SD and SR compared with Ctrl. Selected genes showing state-dependent regulation are indicated. Closed circles represent cortical genes shown in (A), and open circles indicate Atg7 expression in subcortical regions. Hypo., hypothalamus; Cere., cerebellum. C. Schematic of neuron-specific Atg7 knockdown in cortical excitatory neurons. D. Representative fluorescence images showing colocalization of tfLC3 puncta with LAMP1 in cortical excitatory neurons, from shCtrl mice after SD and SR. Arrowheads indicate red tfLC3 puncta lacking LAMP1. Scale bar, 10 μm. E. Similar to (D), but from shAtg7 mice. F. Quantification of total tfLC3 abundance and AV acidification ratio in shCtrl and shAtg7 neurons during SD and SR. Each dot represents data from one mouse (N = 3 mice/group). G. Quantification of LAMP1⁺ and LAMP1⁻ neutral and acidified tfLC3 puncta in shCtrl and shAtg7 neurons during SD and SR. H. Relative proportions of LAMP1⁺ and LAMP1⁻ acidified tfLC3 puncta in shCtrl and shAtg7 neurons after SD and SR. Data are mean ± SEM. One way ANOVA with Bonferroni corrections was used for comparisons between SD and SR in (A), and one sample two-tail t tests were used comparisons against Ctrl reference value of 1 (#P<0.05); Two-way ANOVA followed by Bonferroni correction was used in (F–H) to compare SD versus SR within each genotype and shCtrl versus shAtg7 within SR (##P < 0.01). *P<0.05; **P<0.01; ***P<0.001.

We therefore knocked down Atg7 in cortical excitatory neurons of *CaMKIIα^Cre^*;tfLC3 double transgenic mice using AAV-DIO-shAtg7, with AAV-DIO-scramble-shRNA as the control (shCtrl; **Figure 3C; Supplementary Figure 4A,B**). As expected, SR markedly increased total tfLC3 puncta and AV acidification in shCtrl neurons, whereas both responses were attenuated after Atg7 knockdown (**Figure 3D-F**), indicating a critical role of ATG7 in sleep-dependent autophagy.

To distinguish impaired AP-lysosome fusion from defective acidification after fusion [52], we examined tfLC3 colocalization with lysosomal marker LAMP1 (**Figure 3D-H**). Neutral tfLC3 puncta were predominantly LAMP1⁺ across groups (**Figure 3G**), suggesting preserved efficient AP-lysosome fusion. Unexpectedly, however, we found that more than 40% of acidified puncta in shCtrl neurons during SR lacked LAMP1 (**Figure 3D,H**), indicating a substantial LAMP1⁻ acidified AV population. This population also significantly increased from SD to SR (**Figure 3G,H**). A similar increase was also observed in tfLC3 mice without viral manipulation, excluding a viral expression artifact [53] (**Supplementary Figure 4C-E**). Importantly, while Atg7 knockdown reduced the LAMP1⁺ population in SR, as compared to shCtrl (P=0.052), it almost completely abolished the SR-induced increase in LAMP1⁻ acidified AVs (**Figure 3E,G,H**).

We next explored the identity of the LAMP1⁻ population. Recent studies have shown that LC3 can participate in noncanonical vesicular trafficking and secretion [54, 55]. We therefore examined CD63 (also known as LAMP3), a tetraspanin commonly associated with extracellular vesicles (EVs) as well as lysosomal membranes [55, 56]. Consistent with its dual localization [56], we indeed observed that essentially all LAMP1⁺ tfLC3 puncta were CD63⁺ (**Supplementary Figure 5A,B**). Strikingly, nearly all LAMP1⁻ acidified tfLC3 puncta were also CD63⁺, whereas CD63⁻ puncta were rare (<2%; **Supplementary Figure 5A,B**). The LAMP1⁻ CD63⁺ population increased markedly during SR, from 0.69 to 8.62 puncta/neuron (**Supplementary Figure 5B**). Consistently, SR also increased neuron-derived EVs, as identified by immunogold labeling of EVs containing neuron-selectively expressed CD63-HA and autophagic cargo p62 [57, 58] (**Supplementary Figure 5C-H**). These findings suggest that the sleep-induced LAMP1⁻ population likely represents non-canonical secretory AVs. Because Atg7 knockdown strongly suppressed this population (**Figure 3D-H**), ATG7 appears to support both degradative and putative secretory AVs induced by sleep (**Supplementary Figure 5I**).

### Low neuronal activity facilitates cortical autophagic flux during sleep

In addition to molecular regulation of the autophagy machinery, sleep and wake are accompanied by marked differences in cortical activity [59, 60], while neuronal activity itself has been shown to regulate autophagy in *in vitro* studies [61, 62]. We therefore asked whether neuronal activity contributes to state-dependent changes in autophagy. We examined cFos expression, a marker of recent neuronal activation [63, 64], together with tfLC3 signals in the same cortical neurons from SD or SR mice (**Figure 4A,B**). During SR, we found that cFos-low neurons contained substantially more tfLC3 puncta than neighboring cFos-high neurons, whereas this difference was not apparent after SD (**Figure 4A-D**). Consistently, both tfLC3 puncta abundance and AV acidification were inversely correlated with cFos intensity during SR across all four mice examined, but not during SD (**Figure 4E,F**). Notably, however, both cFos-low and cFos-high neurons showed higher tfLC3 puncta abundance and acidification during SR than SD (**Figure 4C,D**). Thus, while sleep state remained critical for autophagic flux, lower neuronal activity was associated with greater autophagy during sleep.

**Figure 4.**
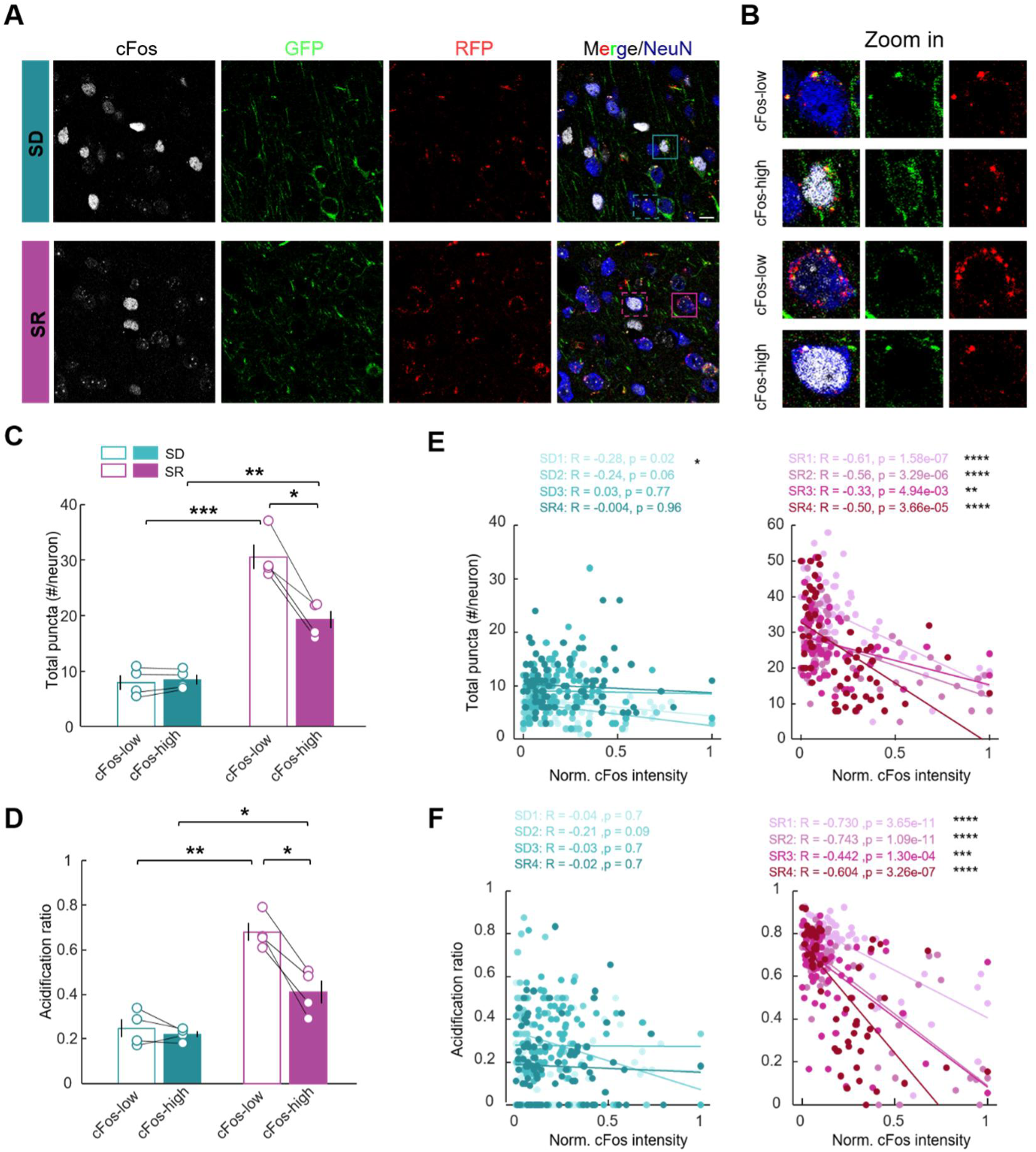
Autophagic flux during recovery sleep inversely correlates with neuronal activity. A. Representative fluorescence images showing tfLC3 and cFos immunostaining in cortical neurons after SD and SR. Boxes indicate neurons shown at higher magnification. Scale bar, 10 μm. B. Higher-magnification views of representative cFos-low and cFos-high neurons from (A). Note that cFos-low neurons in SR, but not SD, showed substantially higher tfLC3 signals. C. Quantification of total tfLC3 puncta in cFos-low and cFos-high neurons after SD and SR. Lines connect values from the same mouse (N = 4 mice/group, >=2 images analyzed for each animal). D. Similar to (C), but for AV acidification ratio. E. Correlation between total tfLC3 puncta and normalized cFos intensity across individual neurons after SD (left) and SR (right). Each dot represents one neuron and colors distinguish individual mice (n>40 neurons for each mice). Lines show linear fits for each mouse; Pearson’s correlation coefficients (R) and P values are indicated. F. Similar to (E), but for AV acidification ratio. Data are presented as mean ± SEM. Two-way mixed ANOVA followed by Bonferroni correction was used in (C, D) to compare cFos-low versus cFos-high neurons within SD and SR, and SD versus SR within the cFos-low and cFos-high groups. *P < 0.05; **P < 0.01; ***P < 0.001; ****P < 0.0001.

To test this relationship directly, we performed chemogenetic activation [65]. We expressed the excitatory DREADD receptor hM3Dq (AAV-CaMKIIα-hM3Dq-BFP) in visual cortical neurons (**Figure 5A**). Mice received either clozapine-N-oxide (CNO, 1.5 mg/kg, i.p.) or saline prior to SR onset (**Figure 5A; Supplementary Figure 6A-C**). Chemogenetic activation modestly reduced NREM sleep (P = 0.055; **Figure 5B,C**), but autophagy remained unchanged in control regions lacking hM3Dq expression (**Figure 5E; Supplementary Figure 6D**). In contrast, hM3Dq-expressing hyperactive neurons showed a significant reduction in both AV abundance and acidification, involving both LAMP1⁺ and LAMP1⁻ acidified AV populations (**Figure 5F,G; Supplementary Figure 6E**). Consistently, these neurons also showed increased phosphorylated S6 (pS6) and p62 levels (**Supplementary Figure 6B,C**), further supporting impaired autophagic clearance [66]. Moreover, this impairment was accompanied by increased intraneuronal Aβ accumulation, with CNO-treated mice showing nearly twice as many Aβ40 puncta (**Figure 5F,H**). In saline-treated mice, neuronal Aβ40 signal was predominantly localized to LAMP1⁺ yellow tfLC3 puncta (**Figure 5F,I**), consistent with accumulation within poorly acidified autolysosomes (paALs) [30, 52]. CNO treatment further increased Aβ40 within paALs (**Figure 5I**), indicating accelerated autophagic Aβ accumulation.

**Figure 5:**
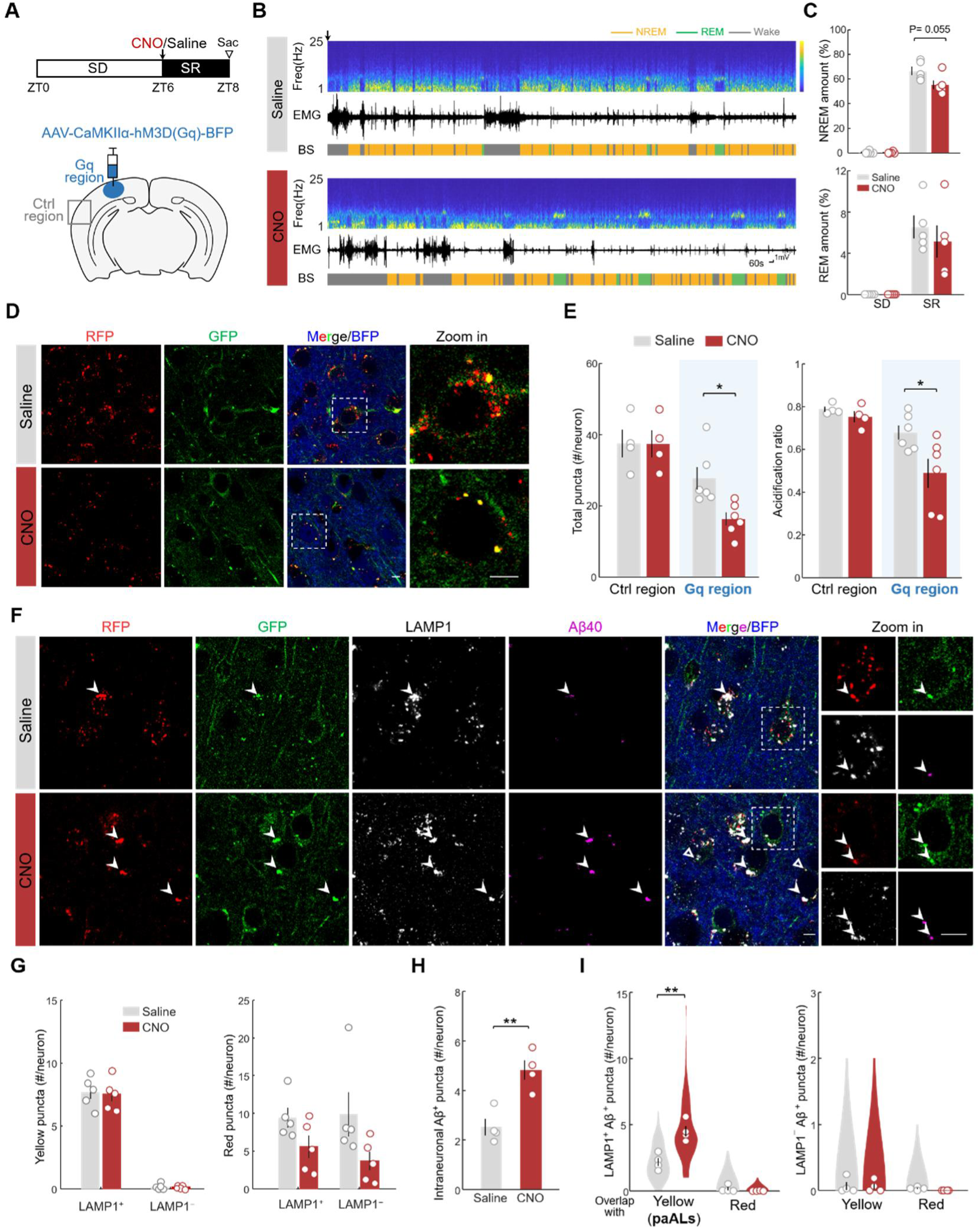
Hyperactivity impairs SR-induced neuronal autophagy and promotes autophagic Aβ accumulation. A. Experimental timeline (top) and viral expression (bottom) for chemogenetic activation of cortical neurons during SR. B. Representative recordings during SR showing EEG spectrogram, EMG trace, and brain states (BS) following intraperitoneal injection of saline or CNO. C. Quantification of NREM sleep (top) and REM sleep (bottom) amount during the 6-h SD and 2-h SR. Each dot represent data from one mouse (N=5 mice/group). D. Representative fluorescence images of tfLC3 puncta in Gq-BFP-expressing cortical neurons, from saline- or CNO-treated mice. Insets show zoom-in views. Scale bar, 5 μm. E. Quantification of total tfLC3 puncta and acidification ratio in Gq-expressing and control cortical regions. Each dot represents one mouse (Ctrl: N=4; Gq: N=6). Blue shading indicates Gq-expressing neurons. F. Representative five-color fluorescence images showing colocalization of tfLC3 with LAMP1 and Aβ40 in Gq-BFP expressing cortical regions. Insets show zoom-in views of individual neurons. Arrowhead: representative LAMP1+ Aβ+ yellow puncta. Scale bar, 5 μm. G. Quantification of LAMP1⁺ and LAMP1⁻ tfLC3 puncta, shown separately for yellow (neutral) puncta (left) and red (acidified) puncta (right), in Gq-expressing cortical regions (N = 5 mice/group). PaALs: poorly acidified autolysosomes. H. Quantification of Aβ40 puncta per neuron in saline- or CNO-treated mice, in Gq-expressing cortical regions (N=4 mice/group). I. Quantification of Aβ40 puncta colocalized with LAMP1⁺ tfLC3 (left) or LAMP1⁻ tfLC3 (right). Note that CNO-mediated hyperactivity during SR induces Aβ accumulation in yellow puncta, i.e., poorly acidified autolysosomes (paALs). Data are presented as mean ± SEM. Unpaired two-tailed *t* test in (C, H). Two-way ANOVA followed by Bonferroni correction was used in (E, G, J) to compare differences between saline and CNO within each condition. * *P < 0.05*; ** *P < 0.01*.

Together, these results indicate that sleep enables ATG7-dependent cortical autophagy, with neuronal quiescence further facilitating efficient clearance and hyperactivity disrupting this process.

### Sleep-specific lysosomal reacidification attenuates Aβ pathology in the cortex of AD mice

We reasoned that if autophagic degradation is most active during sleep, then enhancing lysosomal efficiency specifically during this state should be more effective. To test this hypothesis, we targeted lysosomal acidification, the bottleneck of autophagic clearance in 5xFAD mice [52, 67, 68]. To achieve brain-state-specific manipulation of autophagic flux, we bilaterally expressed AAV-hSyn-lyso-pHoenix, an optogenetic tool for enhancing lysosomal acidification [69, 70], in cortical neurons of 5xFAD mice (∼3-4 months old; **Figure 6A**). Yellow light activates the proton pump (Arch) in lyso-pHoenix, thereby lowering lysosomal pH, as confirmed by *in vivo* imaging (**Supplementary Figure 7A,B**). Using a real-time closed-loop system, light was delivered to one hemisphere selectively during either sleep (N=6 mice) or wakefulness (N=5 mice) over three days, with the contralateral hemisphere serving as an unstimulated control to account for inter-animal variability in Aβ pathology [71] (**Figure 6A,B; Supplementary Figure 7C–F**). To control for nonspecific effects of light, we used a separate cohort expressing lyso-pHluorin without Arch as a light only control (N=4 mice; **Figure 6A**).

**Figure 6.**
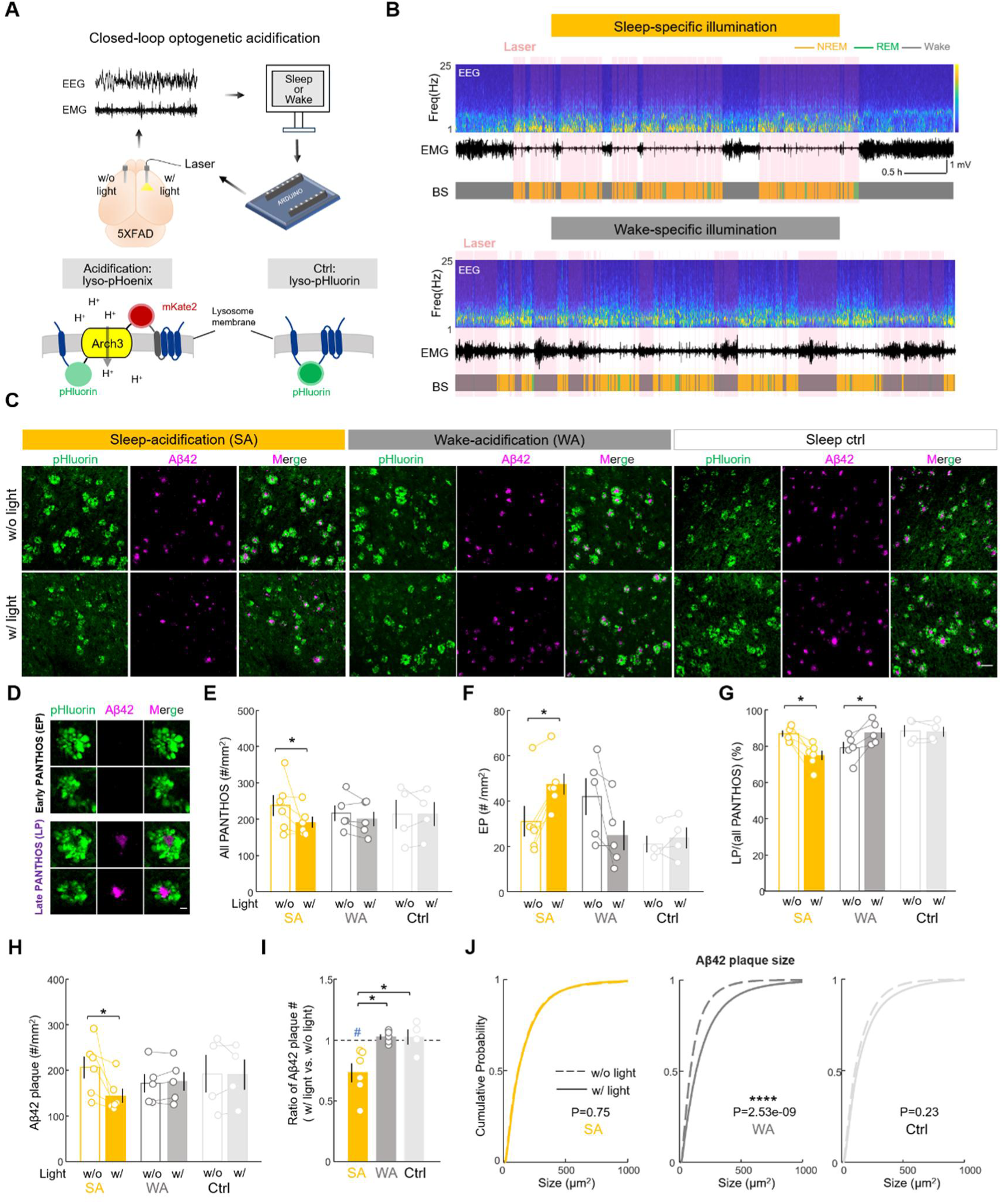
Closed-loop lysosomal acidification during sleep in cortical neurons mitigates Aβ burden in 5xFAD mice. A. Schematic of brain state-specific optogenetic enhancement of lysosomal acidification in 5xFAD mice using Lyso-pHoenix. The contralateral non-illuminated cortex served as an internal control. Lyso-pHluorin, which lacks the light-sensitive proton pump, was used as the light control. B. Representative recordings showing closed-loop light delivery selectively during sleep (top) or wake (bottom). Traces show EEG spectrogram, EMG, brain state (BS), and laser illumination periods (pink shading). C. Representative images of pHluorin and Aβ42 in hemispheres with (w/) or without (w/o) light illumination in sleep-acidification (SA), wake-acidification (WA), and sleep control (Ctrl) groups. Scale bar, 50 μm. D. Representative images of early PANTHOS (EP; without an Aβ42 plaque) and late PANTHOS (LP; with an Aβ42 plaque). Scale bar, 10 μm. E. Quantification of total PANTHOS density in cortical hemispheres w/ or w/o light illumination. Each line represents data from the same mouse (SA, N = 6 mice; WA, N = 5 mice; Ctrl, N = 4 mice). F-H. Similar to (E), but for early PANTHOS density (F), percentage of late PANTHOS (G) and Aβ42 plaque density (H). I. Ratio of Aβ42 plaque density in the hemisphere w/ light relative to the contralateral hemisphere w/o light. Dashed line, ratio = 1; #, significant deviation from 1. J. Cumulative distributions of Aβ42 plaque size in cortical hemispheres w/ and w/o light. Note that wake-specific acidification significantly increased plaque size. Data are presented as mean ± SEM. Two-way mixed ANOVA followed by Bonferroni correction was used in (E-H); one-way ANOVA followed by Bonferroni correction was used for between-group comparisons in (I); one-sample two-tailed t-tests were used for comparisons against 1 in (I); Kolmogorov–Smirnov tests were used in (J). */#P < 0.05; ****P < 0.0001.

Importantly, lyso-pHoenix and lyso-pHluorin also label PANTHOS (poisonous anthos), characteristic flower-like autophagic structures associated with defective acidification and amyloid plaque formation [52, 72] (**Figure 6C; Supplementary Figure 8A,B**), providing an additional readout of neuronal pathology. Consistent with previous finding [52], we found that the majority of PANTHOS were also Aβ42-positive (∼84%, **Figure 6D; Supplementary Figure 9A-C)**. Aβ-negative PANTHOS were smaller and likely represented an earlier pathological stage, whereas Aβ-positive PANTHOS represented a later amyloidogenic stage (**Figure 6D, Supplementary Figure 9D**). We therefore classified them as early PANTHOS (EP) and late PANTHOS (LP), respectively.

We found that PANTHOS abundance was significantly reduced in the illuminated hemisphere, in the sleep-specific lysosomal acidification group, but not in the wake-acidification or light control groups (P<0.05; two-way mixed ANOVA with Bonferroni correction; **Figure 6C,E**; **Supplementary Figure 9E,F**). Sleep-specific acidification also increased the density of EP, while reducing the proportion of late PANTHOS (**Figure 6F,G**), suggesting slower progression toward the amyloidogenic state. Consistently, Aβ42 plaque density was reduced by ∼27% (**Figure 6H,I**). In contrast, wake-acidification increased the proportion of late PANTHOS and significantly enlarged Aβ plaque size (**Figure 6G,J**). These results demonstrate that the effect of enhancing lysosomal acidification depends critically on brain state, with beneficial effects when aligned with sleep.

Taken together, our findings identify sleep as a physiological state that favors efficient neuronal autophagic clearance in the cortex (**Supplementary Figure 9G**). The state-dependent effects of lysosomal acidification further highlight brain-state timing as an important consideration for enhancing intracellular clearance in Alzheimer’s disease.

## Discussion

The fundamental question of why animals sleep has endured for centuries. It is becoming clear that sleep does not serve a single function [73, 74]. Sleep is now recognized to widely support memory consolidation, synaptic remodeling, and metabolic homeostasis [2, 8, 11, 75] , yet the cellular mechanisms that implement these functions remain largely unclear. Autophagy has been implicated in sleep-dependent restoration, but previous findings have been inconsistent. In Drosophila, neuronal AP abundance varies with both circadian phase and prior wake history [26, 27], whereas mammalian studies have reported both increased and decreased autophagy-related markers after sleep loss [28]. By measuring autophagic flux across brain regions and cell types, we show that sleep selectively promotes cortical neuronal autophagy (**Figure 1**), supporting enhanced intracellular clearance during sleep. Our results further suggest that Atg7 upregulation in the cortex during sleep contributes to sleep-associated autophagy, while low neuronal activity further facilitates efficient flux (**Supplementary Figure 9G**). This may be particularly relevant to cortical excitatory neurons, where elevated activity during wakefulness may limit intracellular turnover and increase susceptibility to cellular stress [10, 76, 77].

Time-course analysis showed that cortical autophagy is recruited within tens of minutes (**Supplementary Figure 1E**), and remains elevated during sleep, suggesting that intracellular clearance may represent a rapid and sustained sleep-dependent process [74]. Because REM sleep is largely absent during the initial phase of recovery sleep (**Supplementary Figure 1A**), this increase is likely associated primarily with NREM sleep. Moreover, given the inverse relationship between autophagy and neuronal activity, the lower cortical activity during NREM may in principle permit more efficient autophagic flux compared with REM sleep [59, 60]. Future studies developing brighter reporters for real-time *in vivo* imaging will be needed to investigate autophagic dynamics across NREM and REM sleep, as well as their correlations with specific physiological events such as slow waves, or with local sleep during wakefulness [22]. Our findings also suggest an ATG7-dependent noncanonical route involving CD63-labeled compartments during sleep (**Figure 3 and Supplementary Figure 5)**. Although the secretory fate remains to be validated, it raises the possibility that intracellular degradation and secretion may operate in parallel to remove intracellular cargo from neurons [11, 12, 55, 58, 78]. Beyond bulk intracellular clearance, the preferential targeting of synaptic proteins during sleep suggests that enhanced autophagic flux may also contribute to other sleep-dependent processes such as synaptic remodeling [8, 46] (**Figure 2**), a possibility that warrants further investigation.

Finally, our findings of state-dependent intracellular clearance may have broader implications for neurodegeneration. Sleep disruption, neuronal hyperactivity, and impaired autophagic clearance often coexist in neurodegenerative disease, forming a potential vicious cycle [2, 18, 49, 50, 52, 54, 79–82]. We showed that neuronal activation during sleep reduced autophagy and increased intracellular Aβ, whereas lysosomal acidification had opposite effects across brain states (**Figures 5-6**). Its benefit during sleep is consistent with the concurrent increase in autophagic flux and cargo delivery, while the detrimental effect during wakefulness remains mechanistically unresolved. One possible explanation is that wakefulness is associated with elevated neuronal activity, which can increase APP endocytosis [83], while acidic endolysosomal pH could facilitate BACE1- and γ-secretase-mediated amyloidogenic processing [84, 85]. Thus, enhancing acidification during wakefulness may increase Aβ production and, if not accompanied by a corresponding increase in hydrolytic capacity [86], exacerbate Aβ accumulation.

Such state dependence may also be one factor underlying the variable effects of autophagy-promoting interventions such as rapamycin [87], with treatment timing relative to sleep-wake state as a potential consideration. Recent evidence that the sleep aid lemborexant improves sleep and mitigates tau pathology [88] further suggests that restoring sleep in diseases may benefit proteostasis. Future studies could test whether combining sleep-promoting agents with autophagy-lysosome enhancers improves sleep-autophagy alignment and clearance efficiency, and whether such timing principles translate to clinical intervention. Taken together, our findings support sleep as an important window for neuronal clearance and reveal brain-state timing as a potential determinant of the efficacy of proteostatic interventions in neurodegenerative disease.

## Acknowledgments

We thank L.K. for preparing transgenic mice; Margaret S. Ho for valuable discussions and help with AV identification in EM images; Wei Huang and Yi Zhou for support with western blotting and AV isolation; Chenyan Ma and Peng Yuan for providing the LAMP1 antibody; Qian Hu and Yumei Zhang for assistance with imaging and analysis; Yu Kong and Xu Wang for help with EM; Min Zhang and Songlin Qian for guidance on ultracentrifugation; Jianlong Li, Yue Yin and Qingrun Li for MS sample preparation and data processing. This work was supported by the National Science and Technology Innovation 2030 Major Projects of China (STI2030-2022ZD0206100), CAS Project for Young Scientists in Basic Research (YSBR-106), National Natural Science Foundation of China grants (3237070240 and 8234101126), Space Medical Experiment Program (HYZHXMN01003) and the Benyuan charity foundation.

## Author Contributions

L.Y. performed most experiments and data analysis; Z.X. performed part of the tfLC3 experiments; J.L. performed part of viral injections and data analysis; X.L. assisted with sample preparation for AV proteomics; L.Y. and D.L. wrote the manuscript; D.L. supervised all aspects of the project.

## Data and code availability

The datasets generated during and/or analysed during the current study are available from the corresponding author on reasonable request.

## Materials and methods

### Experimental animals

All animal procedures were approved by the Animal Care and Use Committees of the Center for Excellence in Brain Science and Intelligence Technology (CEBSIT), Chinese Academy of Sciences. Experiments were conducted on adult male and female mice (2–6 months old), including CAG-tfLC3 (CAG-RFP-EGFP-LC3; the Jackson Laboratory, # 027139) [31], *CaMKII^Cre^* (JAX #005359), 5xFAD (JAX #034848) [68], and wild-type C57BL/6J mice (Lingchang Biotechnology Co., Ltd). tfLC3 mice crossed with *CaMKII^Cre^* were used for viral-mediated labeling of excitatory cortical neurons, and tfLC3 mice crossed with 5xFAD were used to visualize PANTHOS structures. Mice were housed under controlled temperature (22– 25 °C) and humidity (50–60%) on a 12-hour light/dark cycle, with ad libitum access to food and water. Animals were randomly assigned to experimental and control groups.

### Surgical procedures

Adult mice (2–4 months old) were anesthetized with 1.5–2% isoflurane and positioned in a stereotaxic frame. Body temperature was maintained at 37.2 °C using a heating pad throughout the procedure. After asepsis, the skin was incised to expose the skull, and the overlying connective tissue was removed.

### For viral injection

A craniotomy (0.5-1 mm in diameter) was made above target brain regions. Stereotaxic coordinates for visual cortex: anteroposterior (AP) -3.0 mm, mediolateral (ML) 2.25 mm, dorsoventral (DV) 1 mm; for motor cortex: AP: +2.46 mm, ML 1 mm, DV 1 mm (from bregma). AAVs (0.3–0.5 µl) were injected using a glass micropipette connected to a Nanoject III (Drummond Scientific). The pipette was positioned 0.05 mm below the target site, then retracted by 0.05 mm before infusion. Virus was delivered at 10-30 nl/min, and the pipette was held in place for 10 min post-injection before slow withdrawal. After suturing, mice recovered on a heating pad before returning to free movement. After surgery, mice were allowed to recover for 2-3 weeks before subsequent experiments. Injection sites were verified post hoc via fluorescence imaging, and animals lacking viral expression were excluded from analysis.

AAVs used in this study included: AAV-EF1α-DIO-GFP-LC3-RFP (modified from Addgene plasmid #84573; gift from Noboru Mizushima) [33]; AAV-EF1α-DIO-hCD63-HA; AAV-CaMKIIα-hM3D(Gq)-BFP; AAV-CaMKIIα-BFP; AAV-EF1α-DIO-shATG7-BFP; AAV-EF1α-DIO-shScramble-BFP; AAV-hSyn-lyso-pHoenix and AAV-hSyn-lyso-pHluorin (constructs modified addgene plasmid #70112 and #70113; gifts from Christian Rosenmund) [69]. AAVs were produced by the CEBSIT Gene Editing Facility.

For gene silencing experiments, three AAVs that contained shRNA targeted different segments of ATG7 were mixed [89]. The shRNA sense sequences were: 5’-TTCTGTCACGGTTCGATAATG-3’, 5’-TCTTACCCTGCTCCATCAAGA-3’, 5’-GCCAACATCCCTGGATACAAG-3’. The scramble sequence for shCtrl was: 5’-CAACAAGATGAAGAGCACCAA-3’.

### For EEG and EMG implantation

Stainless steel screws were inserted into the skull over the frontal and parietal cortices for electroencephalogram (EEG) recording, electromyogram (EMG) electrodes were inserted into the neck muscles, and a reference electrode was placed over the right cerebellum. Insulated leads from the EEG and EMG electrodes were soldered to a pin header, which was secured to the skull using dental cement. After surgery, mice were allowed to recover for at least one week before experiments.

### For optic fiber implantation

At least two weeks after viral injection, optic fibers (0.2 mm diameter; Inper) were bilaterally implanted with fiber tips positioned 0.2 mm above the viral injection site. Implants and EEG/EMG electrodes were secured with dental cement. Mice were allowed at least one week to recover before optogenetic experiments.

### For wide-field imaging

At least two weeks after viral injection of lyso-pHoenix, a 3-mm diameter craniotomy was performed over cortical regions with viral expression, and the exposed dura was covered with a custom-made double-layered glass window. The window was sealed with tissue adhesive (3M), and a custom-made stainless-steel headplate with a screw hole was embedded into the dental acrylic, allowing for head-fixed recording. Mice were allowed at least two weeks to recover before imaging.

### EEG/EMG recordings and sleep-wake scoring

Mice were in their home cage placed in a sound-attenuation box. EEG and EMG electrodes were connected to flexible recording cables via a mini-connector. EEG and EMG signals were recorded and amplified with Medusa Server (Bio-signal) or Open Ephys at 1000 Hz.

For offline sleep scoring, spectral analysis was carried out using fast Fourier transform (FFT), and brain states were classified into wake (desynchronized EEG and high EMG activity) and sleep state (NREM: synchronized EEG with high delta power (1-4 Hz) and low EMG activity; REM: high EEG theta power (6-10 Hz) and low EMG activity, with an epoch length of 5 s. Short periods of EEG desynchronization during NREM sleep that lasted for less than 15 s were also considered as NREM sleep. Sleep stages were manually proofed using a custom-written graphical user interface [90].

For online sleep scoring in the closed-loop system, EEG and EMG signals recorded by Medusa Server (Bio-signal) were streamed to MATLAB at 500 Hz. Thresholds for delta power and EMG amplitude were determined from prior recordings in the same animal. Sleep was scored when delta power exceeded its threshold or EMG power fell below its threshold, encompassing both NREM and REM sleep, with an epoch length of 2.5 s.

### Sleep Deprivation and Recovery

Sleep deprivation (SD) began at zeitgeber time 0 (ZT0) and lasted to ZT6, during which mice were kept awake using novel objects or gentle stimulation (e.g., tapping, cage shaking, or light touch). For sleep recovery (SR), mice were allowed ∼2 hours of undisturbed rest (SD6 – SD8) immediately following SD. Control animals were collected at ZT6 or ZT8 to match the circadian time of the corresponding SD or SR group, respectively, to minimize potential circadian confounds. In most experiments, EEG and EMG were continuously recorded to verify wakefulness during SD and sleep rebound during SR.

### Closed-loop optogenetic acidification

Closed-loop stimulation was initiated based on real-time sleep scoring as described above. When two consecutive epochs of the target state were identified, MATLAB triggered an Arduino microcontroller (UNO) to deliver a TTL signal that activated a 583-nm laser (10 Hz, 10 mW, Shanghai Laser). Illumination was unilateral for 3 consecutive days, with the contralateral cortex as internal control. Light was delivered during sleep for sleep-specific acidification and lyso-pHluorin controls, and during wakefulness for wake-specific acidification. To ensure comparable illumination across groups, total light exposure was capped at ∼30 hours per mouse over the 3-day period.

### Wide-field imaging

Wide-field calcium imaging was performed in head-fixed mice expressing lyso-pHoenix. Fluorescence signals were acquired with a stereomicroscope (MVX10, Olympus; ×1 objective) coupled to a CMOS camera (Moment; 7 MP; 17.5 mm diagonal; 50 fps) through a 535/45 nm emission filter (Chroma). Excitation light was delivered by a CoolLED pE-300 Ultra system, with frame-synchronized alternation between 470 nm (for pHluorin excitation) and 410 nm (for hemodynamic artifact correction), controlled by a custom setup triggered by the camera. Images were captured using Micro-Manager software at a resolution of 1,024 × 1,024 pixels (2× binning, 16-bit depth) with an exposure time of 50 ms per frame. During imaging, 583 nm laser (30 mW, ∼30-70s each trial) was delivered through the cranial window at randomly timed intervals ranging from 25 to 80 seconds. Each session consists of 2-4 trials.

### Drug infusion

For chemogenetic activation, mice expressing CaMKIIα-hM3Dq received intraperitoneal injections of clozapine-N-oxide (CNO; 1.5 mg/kg body weight) or saline at SR onset (ZT 6). EEG and EMG were continuously recorded, covering 6-h SD followed by 2-h SR before sacrifice for tissue collection.

### Western blotting

Mice were anesthetized with isoflurane, decapitated to collect fresh brain tissue. Targeted cortical regions were lysed in pre-chilled RIPA buffer containing 50 mM Tris-HCl (pH 7.4), 150 mM NaCl, 1% Triton X-100, 1% sodium deoxycholate, 0.1% SDS, sodium orthovanadate, sodium fluoride, EDTA, and 1% protease inhibitor cocktail (Thermo Fisher Scientific). Lysates were centrifuged, and supernatants were collected. Protein concentration was determined using the BCA protein assay kit (BeyoBCA). Equal amounts of protein (10 or 20 μg per lane) were separated on 4–20% gradient SDS-PAGE gels (EpiZyme, China) and transferred to 0.22-μm PVDF membranes (Millipore). Membranes were incubated overnight at 4 °C with primary antibodies against anti-Atg7 (1:1000, Abcam, Cat# ab133528) or Mouse anti-β-Actin (1:3000, ABclonal, Cat# AC004, RRID: AB_2737399), followed by incubation with HRP-conjugated secondary antibodies (Beyotime). Blots were developed using an enhanced chemiluminescence kit (Beyotime).

### Quantitative real-time PCR

Mice were anesthetized with isoflurane and transcardially perfused with ice-cold saline to collect fresh brain tissue. Total RNA from targeted cortical tissues was extracted using TRIzol reagent (Invitrogen) and quantified with a NanoDrop 2000 spectrophotometer (Thermo Fisher Scientific). Complementary DNA (cDNA) was synthesized using the PrimeScript RT reagent Kit (TaKaRa) according to the manufacturer’s instructions. Quantitative PCR was performed using TB Green™ Premix Ex Taq (TaKaRa) in 96-well optical plates on a LightCycler system (Roche Diagnostics). mRNA expression levels were normalized to the reference gene Actb and calculated using the 2^–ΔΔCT method. Primer sequences are listed below.

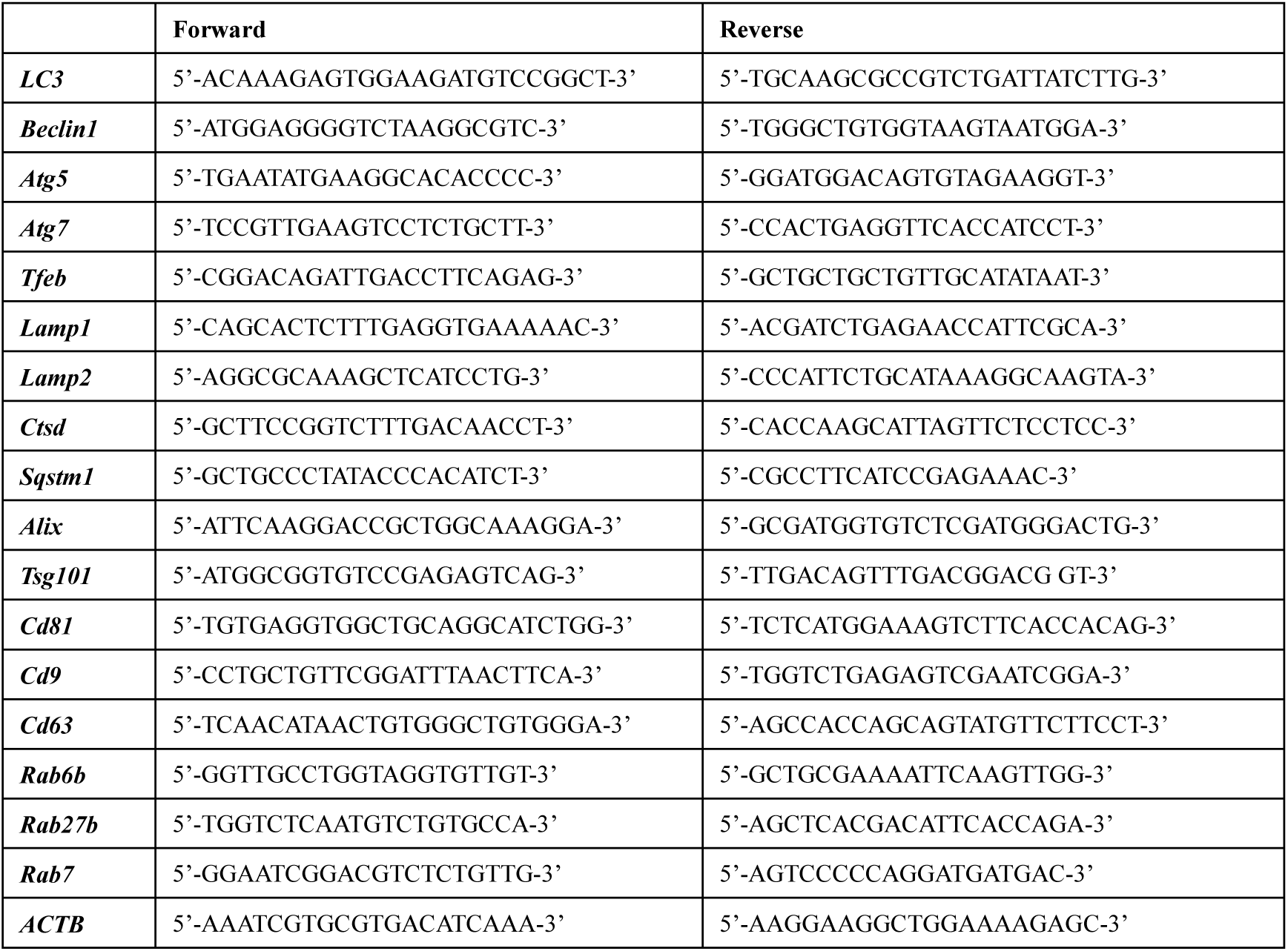

### Histology and immunofluorescence

Mice were anesthetized with isoflurane and transcardially perfused with ice-cold saline followed by 4% paraformaldehyde (PFA) in PBS. Brains were post-fixed in 4% PFA overnight at 4 °C, then cryoprotected in 30% (w/v) sucrose in PBS at 4 °C. Coronal sections (40-50 μm) were obtained using a cryostat (Leica CM1860). Free-floating sections were washed in PBS, permeabilized with 0.3% Triton X-100 for 20 min, and blocked with 5% bovine serum albumin (BSA) for 1 h at room temperature. Sections were then incubated with primary antibodies overnight at 4 °C, followed by PBS washes and a 4-h incubation with fluorescence-conjugated secondary antibodies. After final PBS washes (3 × 10 min), sections were mounted, air-dried, and coverslipped using VECTASHIELD antifade mounting medium (Vector Laboratories). Edges were sealed with nail polish. The following primary antibodies were used:

The following secondary antibodies were used: donkey anti-mouse Alexa Fluor™ 405 (Cat# A48257), Alexa Fluor™ 488 (Cat# A21202) or Alexa Fluor™ 647 (Cat# A-31571); donkey anti-rabbit Alexa Fluor™ 647 (Cat# A-31573); donkey anti-rat Alexa Fluor™ 647 (Cat# A48272TR) (all from Thermo Fisher Scientific); goat anti-rabbit Multi-rAb CoraLite® Plus 750 (Proteintech, Cat# RGAR006).

| **Primary antibodies** | **Source** | **Cat #** |
|---|---|---|
| Mouse anti-NeuN (1:1000) | Millipore | Cat# MAB377 |
| Mouse anti-GFAP (1:500) | Millipore | Cat# MAB360 |
| Rabbit anti-IBA1 (1:500) | Wako Pure Chemical | Cat# 019-19741 |
| Mouse anti-TH (1:1000) | Millipore | Cat# MAB318 |
| Rabbit anti-CC1 (1:500) | Cell Signaling Technology | Cat# CST 89332 |
| Rabbit anti- CaMKII (1:500) | Proteintech | Cat# 13730-1-AP |
| Guinea Pig anti-PV (1:500) | Immunostar | Cat# 24428 |
| Rabbit anti-p62 (1:300) | MBL | Cat# PM045 |
| Rat anti-LAMP1 (1:500) | Developmental Studies Hybridoma Bank | Cat# 1D4B |
| Rabbit anti-LAMP1(1:300) | Abcam | Cat# ab24170 |
| Rabbit anti-HA-Tag (1:500) | Cell Signaling Technology | Cat# CST3724 |
| Mouse anti HA-Tag (1:500) | ABclonal | Cat# AE008 |
| Rabbit anti-c-Fos (1:1000) | Synaptic Systems | Cat# 226 003 |
| Rabbit anti-phospho-S6 (1:500) | Invitrogen | Cat# 44-923G |
| Rabbit anti-Aβ,1-40 (1:300) | Millipore | Cat# AB5074P |
| Rabbit anti-Aβ,1-42 (1:300) | Invitrogen | Cat# 700254 |

### Confocal imaging and image analysis

Confocal images were acquired using an Olympus FV3000 and Leica Stellaris 8 STED microscope (for 5-color fluorescence imaging as shown in Fig.5f). Laser power and detector settings were kept constant across samples for each staining condition. For each mouse, at least three images were collected per slide using the following objectives: Plan-Apo 20× (NA 0.75, WD 0.6, dry, Olympus), 60× oil (NA 1.3, WD 0.3, Olympus), and 63× oil (NA 1.4, WD 0.14, Leica), with a resolution of 1024 × 1024 pixels and z-step intervals of 0.5 or 1 μm.

Image analysis was performed using Fiji (ImageJ). For tfLC3 puncta analysis, high-resolution 60× images acquired under identical settings were analyzed from raw 3D image stacks. Because the RFP signal showed a higher signal-to-noise ratio, RFP-positive puncta were identified using a threshold of >3-fold above local background fluorescence. The GFP signal had a lower signal-to-noise ratio and was therefore identified using a >2-fold-above-background threshold. Puncta were manually identified based on intensity and discrete morphology rather than a fixed size threshold. For colocalization of multi-channel signals, those with centroid distances within 2 pixels were considered colocalized. For fluorescence intensity quantification of p62, GFP-LC3-RFP, and pS6, z-stacks were converted to maximum intensity projections, and mean intensities within NeuN^+^ neurons were measured. For PANTHOS and Aβ plaque analyses, a customized Fiji macro was used for ROI segmentation and size quantification.

### Isolation and purification of LC3^+^ autophagic vesicles from mouse brain

AVs were purified using a modified protocol combining previously published methods [34–37], to enrich membrane-protected cargo from LC3^+^ AVs, including both APs and ALs. Briefly, forebrains from mice, following SD or SR were rapidly dissected and homogenized in Buffer A containing 10% (w/v) sucrose, 10 mM HEPES, and 1 mM EDTA, pH 7.3, using a Dounce homogenizer. The homogenate was diluted in Buffer B containing 250 mM sucrose, 10 mM HEPES, and 1 mM EDTA, pH 7.3. GPN treatment was omitted to preserve autolysosomes. Samples were incubated at 37°C for 7 min and then centrifuged at 2,000 × g for 2 min at 4°C to remove nuclei and cell debris. The post-nuclear supernatant was layered onto a pre-chilled discontinuous Nycodenz density gradient consisting of 9.5% and 22.5% Nycodenz prepared in Buffer B and ultracentrifuged at 141,000 × g for 1 h at 4°C using a swinging-bucket rotor (Beckman). The AV-enriched interface was collected and further purified by centrifugation through a 33% Percoll / 22.5% Nycodenz gradient at 72,000 × g for 30 min at 4°C. AV fraction at the lower interface was collected and mixed with 0.7 volumes of 60% OptiPrep, overlaid sequentially with 30% OptiPrep and Buffer B, and centrifuged at 71,000 × g for 30 min at 4°C, to remove Percoll silica. The resulting AV-containing pellet was resuspended in 1× DPBS to a final volume twice that of the original pellet. Protein concentration was measured using a BCA assay.

LC3-positive AVs were further enriched from the isolated AV fraction by immunoprecipitation using the Dynabeads™ Protein G Immunoprecipitation Kit (Thermo Fisher Scientific), according to the manufacturer’s instructions with minor modifications. Briefly, 10 μg of anti-LC3 antibody (Sigma-Aldrich, L7543) or control IgG antibody (Sigma-Aldrich, I8140) was incubated with 50 μL of Dynabeads™ Protein G magnetic beads in binding buffer for 30 min at room temperature with gentle rotation. The antibody-conjugated beads were then washed with binding buffer to remove unbound antibody. Equal amounts of AV input were added to the antibody-conjugated beads and incubated for 1 h at room temperature with gentle rotation, to capture LC3+ AVs. The beads were washed by washing buffer to remove unbound material. The bead-bound LC3^+^ AVs were then treated with Proteinase K (20 ng/μL) on ice for 20 min to digest proteins exposed on the outer surface of the vesicles or otherwise externally associated with them. Laemmli buffer was then added to a final concentration of 1×, and samples were heated at 95°C for 5 min to lyse the vesicles. Following magnetic separation, the resulting eluates were collected for proteomics.

### Mass spectrometry and data analysis

Label-free quantitative proteomics was performed following a previously established protocol [91]. Proteins in the eluates were extracted, acetone-precipitated, and denatured in 8 M urea, followed by reduction (5 mM TCEP, Thermo Fisher Scientific and alkylation (10 mM iodoacetamide, Sigma-Aldrich). Samples were digested with trypsin (1:50 w/w, Promega) for 16h at 37°C and desalted with STAGE Tips. An equal amount of peptide (1 μg) from each sample was loaded for LC-MS/MS analysis. Peptides were analyzed on a Q Exactive MS coupled to an Easy-nLC 1200 system (Thermo Fisher). MS was operated in data-dependent mode with full MS scans (m/z 300–1800, resolution 70,000), followed by HCD MS2 scans (resolution 17,500, collision energy 28).

Raw data were processed with MaxQuant (v2.1.4.0) [92] using the UniProt Mus musculus database plus common contaminants. Carbamidomethylation of cysteine was specified as a fixed modification; oxidation (M) and acetylation (protein N-term) as variable. Trypsin/P was used as the enzyme specificity, allowing up to two missed cleavages. The false discovery rate was controlled at 1% at both peptide and protein levels. Label-free quantification was enabled in MaxQuant for relative protein abundance analysis. Intensity-based absolute quantification (iBAQ) values calculated from MaxQuant were used for downstream analysis after removal of reverse sequences and potential contaminants. iBAQ values were median-normalized within each biological replicate. Missing values in IgG control samples were imputed from a normal distribution with a width of 0.3 and a downshift of 1.8, whereas missing values in LC3-IP samples were retained as NA and were not imputed. Proteins identified at least 1 unique peptide in more than two biological replicates were retained for further analysis. Differentially enriched proteins were identified using Student’s t-test, with a nominal P value threshold of < 0.05. Full proteomic datasets are provided in Data S1.

Synaptic proteins and their subcellular localizations were annotated using the SynGO database (https://syngoportal.org/) [93]. Mitochondria proteins and their subcellular localizations were annotated with MitoCarta3.0 database (https://personal.broadinstitute.org/scalvo/MitoCarta3.0/) [94]. Proteins associated with other cellular components, including the endoplasmic reticulum (ER) and lysosome were annotated using GO cellular component terms, and further validated using UniProt (https://www.uniprot.org) and COMPARTMENTS database (http://compartments.jensenlab.org). Gene enrichment analysis was performed using GO biological process terms (via WebGestalt online tool).

### Transmission electron microscopy and quantification

Mice were transcardially perfused with 0.1 M phosphate buffer (PB, pH 7.4, 37°C), followed by 4% paraformaldehyde (PFA) and 1% glutaraldehyde (GA) in PB at 4°C. Brain tissues containing the visual cortex were sectioned into 100-μm slices and post-fixed overnight at 4°C in 4% PFA with 2.5% GA. Sections were washed in PB and deionized water, treated with 1% osmium tetroxide (OsO₄) for 30 min, followed by a single heavy-metal counterstain with 1% aqueous uranyl acetate (UA) overnight. After rinsing, samples were dehydrated in a graded ethanol series and infiltrated with Epon 812 resin (Electron Microscopy Sciences) as follows: 3:1 ethanol: resin for 1 h; 1:1 for 2 h; 1:3 for 2 h; then 100% resin overnight and again for 8 h. Sections were embedded between ACLAR films (Ted Pella), flattened with glass slides, and polymerized at 60°C for 48 h. Chips from the visual cortex were trimmed and mounted on resin blocks for ultrathin sectioning. Sections (∼70 nm) were cut and imaged using a JEM-1230 transmission electron microscope (JEOL). No additional UA or lead citrate staining was performed after ultrathin sectioning, as in our preliminary tests it increased overall electron density and reduced the morphological distinction between ALs and lysosomes. Under this pre-embedding single-staining protocal, membrane contrast and internal cargo morphology were sufficiently preserved for classification of AV structures.

For identification of AV structures, APs were defined as double-membrane vesicles enclosing relatively intact cytoplasmic material or organelles. Mitochondria were distinguished by their characteristic cristae and morphology. ALs were identified as single-membrane compartments containing heterogeneous, partially degraded cargo, typically with lower and more heterogeneous electron density than mature lysosomes. Lysosomes were distinguished by their relatively uniform electron-dense content and absence of recognizable undegraded cargo. MVBs were identified as single-membrane compartments containing multiple intraluminal vesicles. These criteria were applied consistently across all groups. For quantification, APs and ALs were counted from single ultrathin sections through the perinuclear region of neuronal somata. Measurements from more than 40 neuronal soma sections per animal were averaged, with animals used as the statistical unit.

### Isolation of cortex-derived EVs, immunogold-EM and EV immunofluorescence

Fresh visual cortices from hCD63-expressing mice were dissected, with the same amount of starting tissue used for the SD and SR groups. Tissues were enzymatically dissociated in Hibernate A medium containing 20 U/mL papain (Worthington) at 37°C for 20 min. Enzymatic activity was halted by gentle homogenization in twice the volume of ice-cold Hibernate A supplemented with 0.1% protease inhibitor cocktail. The homogenate was filtered through a 40-μm cell strainer (BD Biosciences), and EVs were isolated from the filtrate following a previously described protocol [95]. Briefly, the filtrate was sequentially centrifuged at 300×g for 5 min, 2,000×g for 10 min, and 10,000×g for 30 min at 4°C to remove cells, membranes, and large debris. The supernatant was then passed through a 0.22-μm syringe filter (Millipore) and ultracentrifuged at 100,000×g for 70 min at 4°C. The EV pellet was washed and recentrifuged at 100,000×g for 70 min, then resuspended for TEM and immunofluorescence analyses.

For Immunogold EM of purified EVs, EVs were fixed in 2% PFA for 10 min at RT and adsorbed onto carbon-coated grids for 1 h. Grids were washed three times with PBS, quenched in 50 mM glycine for 10 min, and blocked with 1% BSA for 30 min. Grids were incubated with primary antibodies against Rabbit anti-CD63 (1:30, ABclonal, Cat# A5271, RRID:AB_2766092) or Rabbit anti-HA (1:20, Cell Signaling Technology Cat# 3724, RRID:AB_1549585) overnight at 4°C, washed, re-blocked, and incubated with 5-nm anti-rabbit gold-conjugated secondary antibody (Sigma-Aldrich, Cat# G7277, RRID:AB_259951 ) for 1 h. Samples were post-fixed in 1% GA and negatively stained with 1% UA for 5 min before imaging on a Talos L120C at 80 kV. Note that equal amounts of starting tissue were used for both groups, and paired samples were processed in parallel within the same batch using identical isolation and immuno-EM procedures to minimize variability in sample recovery and processing loss. For quantification of immunogold-EM of EVs, 250 imaging fields were analyzed under identical imaging conditions. Because hCD63-HA was expressed in only a subset of cortical excitatory neurons and HA immunogold labeling was sparse, HA-negative EVs could not be interpreted as CD63-negative EVs or used to define the total EV pool. We therefore counted HA-positive EVs directly across the same total imaged area without normalization to total EV number and compared their absolute abundance between SD and SR.

For fluorescence analysis, EV suspensions were smeared onto poly-L-lysine–coated slides, air-dried, and fixed in 4% PFA for 15 min. After three 10-min washes in PBS, slides were blocked in 5% BSA for 1 h, followed by a 90-min incubation with primary antibodies: Rabbit anti-p62 (1:300, MBL, Cat# PM045) and Mouse anti-HA (1:500, ABclonal, Cat# AE008). Slides were then washed and incubated for 90 min with the following secondary antibodies: donkey anti-mouse Alexa Fluor™ 488 (Thermo Fisher Scientific, Cat# A-21202), donkey anti-rabbit Alexa Fluor™ 594 (Thermo Fisher Scientific, Cat# A-21207). Finally, slides were mounted with antifade mounting medium for fluorescence imaging.

## Statistical analysis

Data are presented as mean ± SEM unless otherwise indicated. No statistical methods were used to predetermine sample size. Comparisons between two groups were performed using paired two-tailed Student’s t-tests for measurements obtained from the same animal, or unpaired two-tailed Student’s t-tests for independent groups, as appropriate. One-sample t-tests were used for comparisons against a reference value. Comparisons involving multiple groups or factors were performed using one-way ANOVA, two-way ANOVA, or two-way mixed ANOVA for matched measures, followed by Bonferroni correction where appropriate. Welch’s one-way ANOVA followed by Games–Howell correction was used in Figure 2E, where group sizes differed substantially. Kolmogorov–Smirnov tests were used for cumulative distributions, Fisher’s exact test for categorical proportions, and Pearson’s correlation for correlation analyses. Exact statistical tests, sample sizes, and significance thresholds are provided in the corresponding figure legends. Statistical analyses were performed using GraphPad Prism (v8.0) or MATLAB (R2019b).

## Supplemental Figures and Figure Legends

**Supplementary Figure 1.**
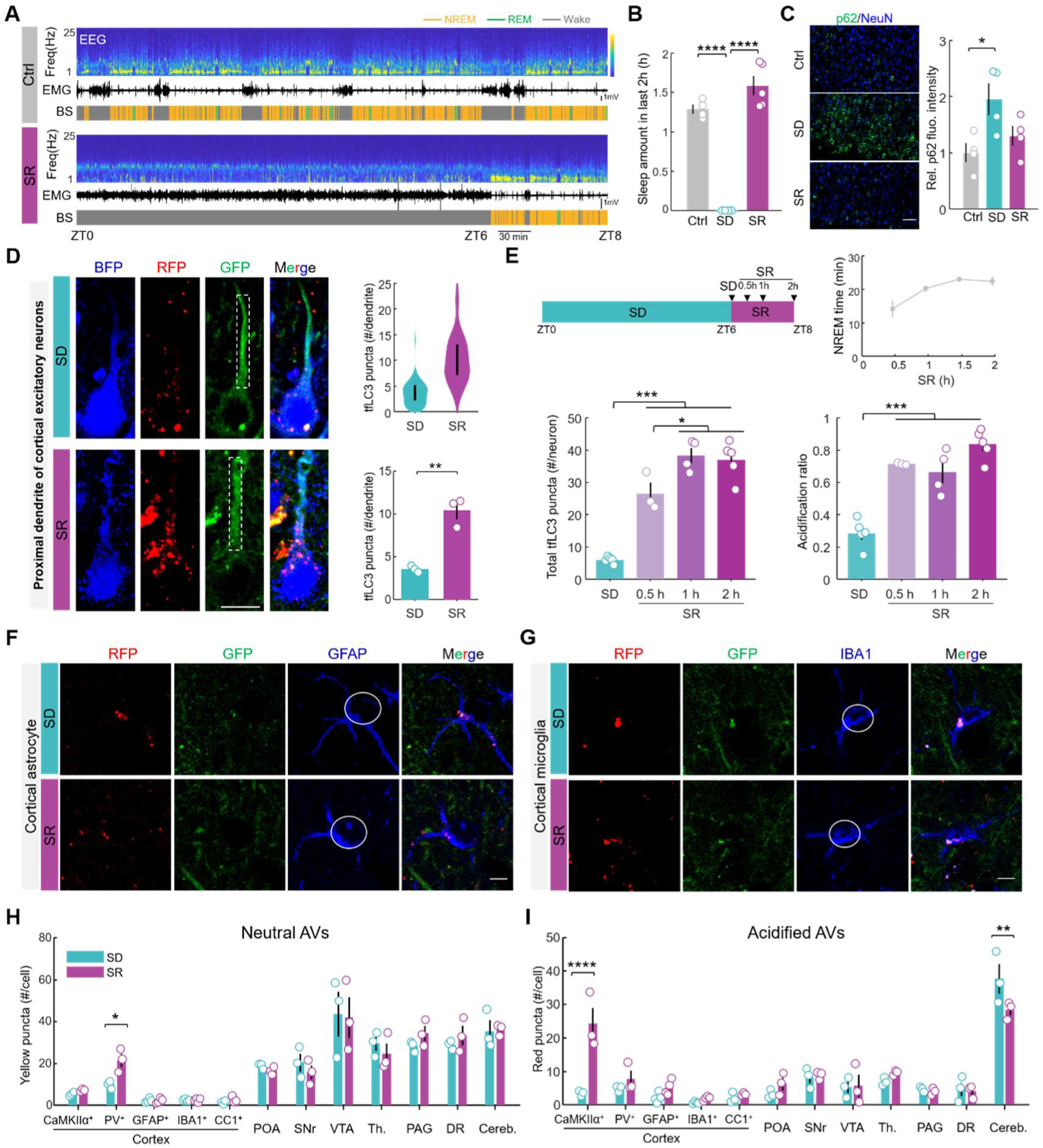
Effects of sleep-wake states on autophagy flux across the brain. A. Representative sleep recordings from Ctrl and SR mice, shown are EEG spectrograms, EMG traces, and brain states (BS). B. Sleep amount during the 2-h periods prior to sacrifice in Ctrl (N= 4 mice), SD (N=4), and SR (N=5) groups. C. Representative fluorescence images (left) and quantification (right) showing neuronal p62 levels across Ctrl, SD and SR (N = 4 mice/group). Scale bar, 50 μm. D. ***Left:*** Representative images and quantification of tfLC3 puncta in proximal apical dendrites of AAV-BFP labeled cortical excitatory neurons under SD and SR conditions. Dendritic segments within ∼30 μm of the soma were analyzed. Dashed rectangle, apical dendrite trunk. Scale bars, 10 μm. ***Right:*** Distribution of tfLC3 puncta numbers across dendrites and group quantification (N = 3 for each Group). E. ***Top:*** Schematic of the time-course analysis during the first 2 h of recovery sleep and time spent asleep in each 30-min bin. **Bottom:** Time-course analysis of cortical neuronal autophagic flux after 0.5, 1, and 2 h of recovery sleep, showing total tfLC3 puncta abundance and acidification ratio. Mice in the 0.5-h SR group slept for ∼15 min before tissue collection. F. Representative fluorescence images of tfLC3 puncta in GFAP⁺ astrocytes. Circles outline somata. Scale bars, 5 μm. G. Similar to (F), but for IBA1^+^ microglia. H. Quantification of neutral tfLC3 puncta abundance across cell populations under SD and SR conditions (N=3 or 4 mice/group). Each dot represents data from one mouse. I. Similar to (H), but for acidified tfLC3 puncta abundance. Error bars represent SEM. Unpaired two-tailed t-test was used in (D); One-way ANOVA with Bonferroni’s correction was used in (B), (C), (E). Two-way ANOVA with Bonferroni’s correction in (H), (I). *P < 0.05; **P <0.01; ***P<0.001; ****P < 0.0001.

**Supplementary Figure 2.**
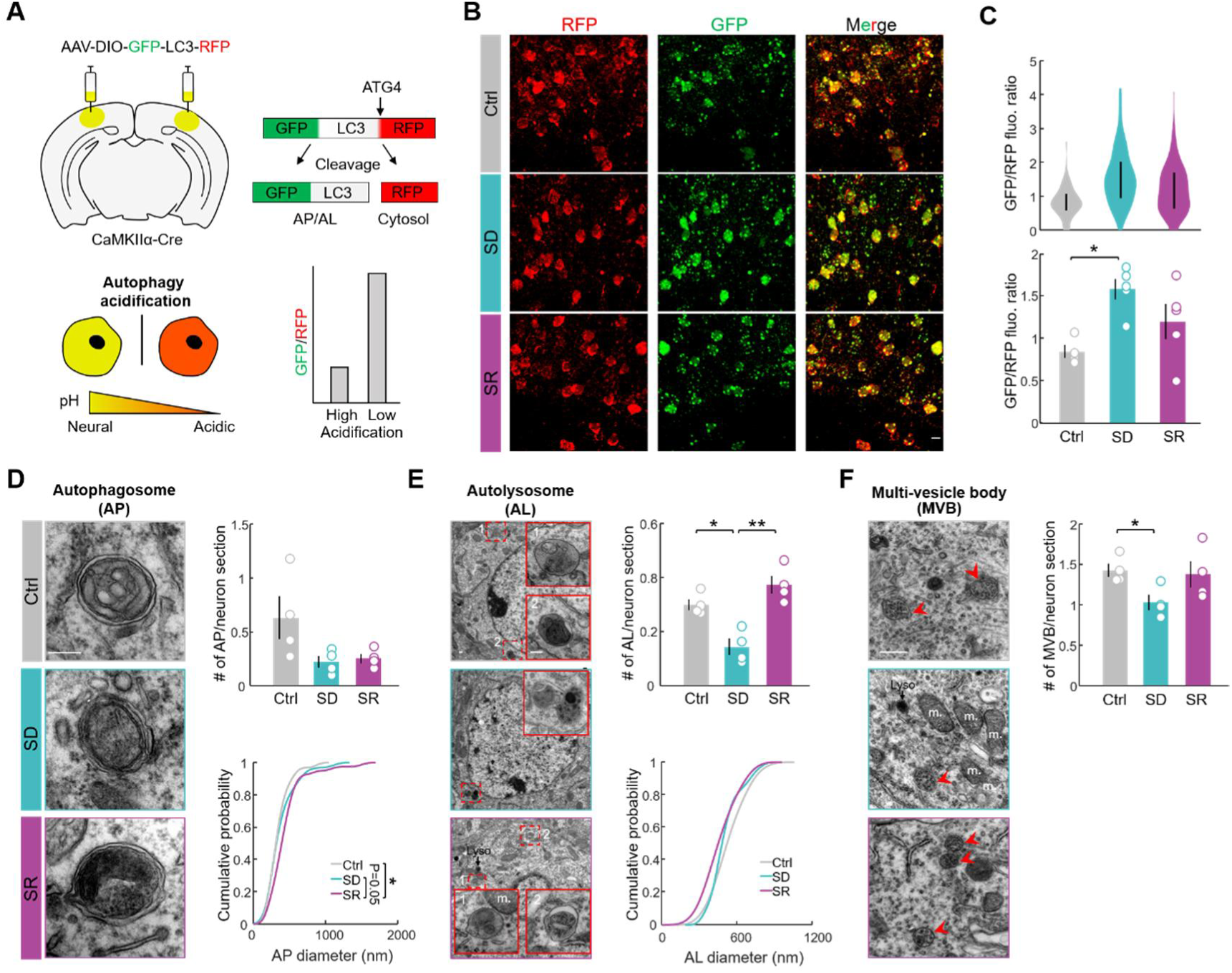
Sleep promotes autophagic acidification and maturation in cortical neurons. A. Schematic of the GFP-LC3-RFP ratiometric reporter and cortical viral injection. Cytosolic RFP serves as an internal control, with a lower GFP/RFP fluorescence ratio indicating greater AV acidification. B. Representative fluorescence images of GFP-LC3-RFP expression in cortical neurons from Ctrl, SD, and SR mice. Scale bars: 10 μm. C. Quantification of the GFP/RFP fluorescence ratio across Ctrl, SD, and SR groups. ***Top:*** distribution of single-cell GFP/RFP ratios (∼110-180 neurons per group) pooled across multiple mice. Vertical bars indicate IQR. Bottom: average GFP/RFP ratio for each mouse (Ctrl, N=4; SD and SR, N=5 each). D. ***Left:*** representative transmission electron microscopy images of autophagosomes (APs) in cortical neurons, identified as double-membrane vesicular structures enclosing cytoplasmic material. ***Right***: quantification of AP abundance and cumulative distribution of AP diameter (N=4 mice per group). Scale bars, 200 nm. E. Similar to (D), but for autolysosomes (ALs), identified as single-membrane vesicular structures containing partially degraded or electron-dense material. Insets show enlarged views of representative ALs. Lyso, lysosome; m., mitochondrial. F. Representative TEM images and quantification of multivesicular bodies (MVBs) in cortical neurons (N = 4 mice/group). Arrowheads indicate MVBs, identified as single-membrane structures containing multiple small intraluminal vesicles. Scale bars, 500 nm. Error bars represent SEM. One-way ANOVA with Bonferroni’s correction was used in (C-F), and Kolmogorov–Smirnov test in cumulative distributions in (D) and (E) *P < 0.05; **P <0.01; ***P<0.001; ****P < 0.0001.

**Supplementary Figure 3.**
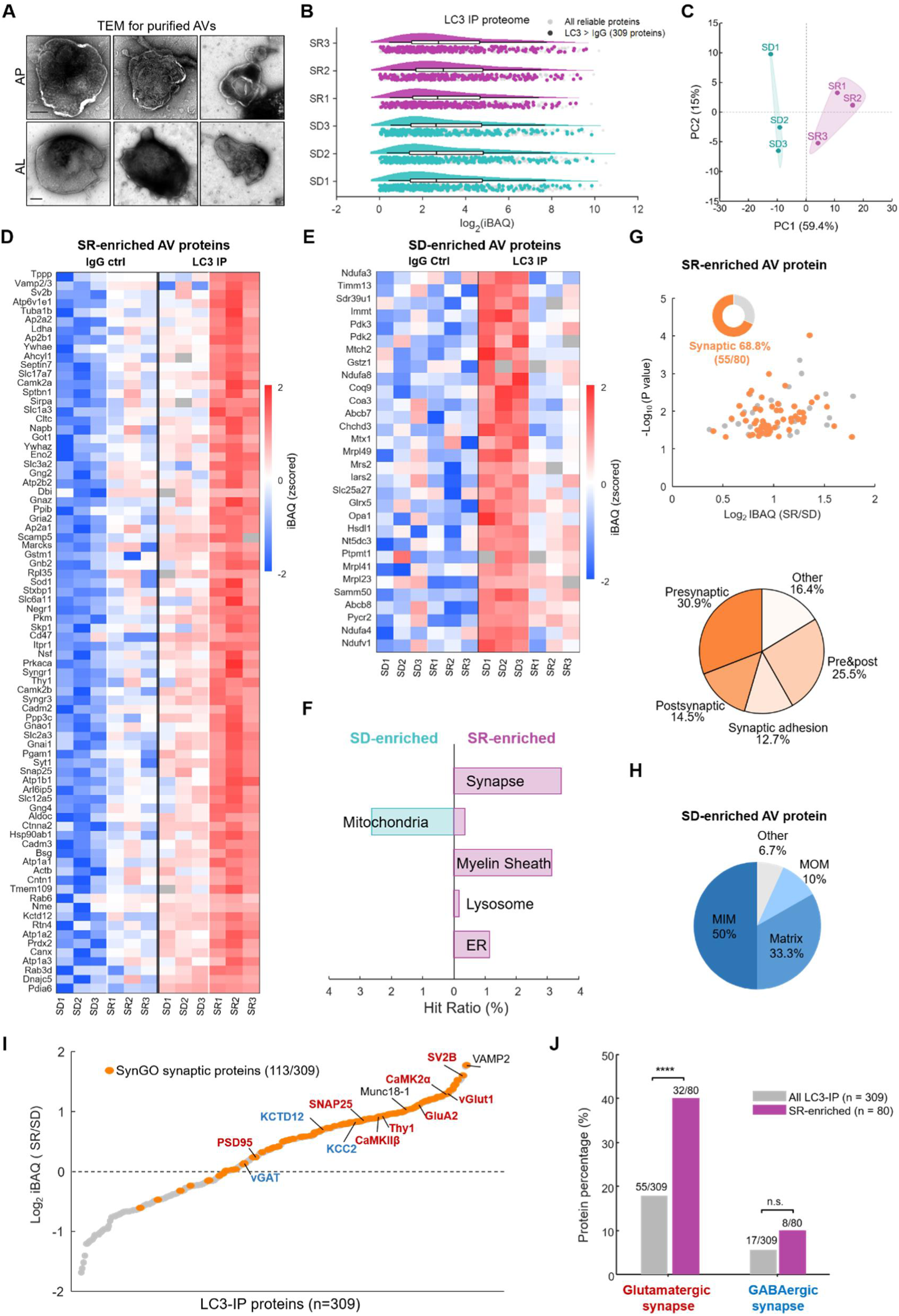
Selective targeting of excitatory synaptic components by AVs during sleep. A. Representative TEM images of isolated cortical APs and ALs. Scale bars, 200 nm. B. Distribution of protein abundance across individual SD and SR samples (N = 3 biological replicates per group). C. Principal-component analysis (PCA) of AV proteomes from SD and SR samples. Each point represents one biological replicate. D. Heatmap showing the abundance of SR-enriched AV proteins (n=80) across individual SD and SR mice, in IgG controls and LC3-IP samples. Values are normalized by protein. E. Similar to (D), but for 30 SD-enriched AV proteins. F. Hit ratio of major subcellular components among SD- and SR-enriched AV proteins. G. **Top:** volcano plot of SR-enriched proteins, with synaptic proteins highlighted (n=55). **Bottom:** pie chart showing the subcellular localization of these synaptic proteins. H. Pie chart showing subcellular localization of SD-enriched mitochondrial proteins. I. Ranked log₂ iBAQ(SR/SD) values for all 309 AV proteins, with synaptic proteins highlighted in orange. Selected excitatory synaptic components are labeled in red and inhibitory components in blue. J. Relative proportions of glutamatergic and GABAergic proteins among all AV synaptic proteins (gray) and SR-enriched synaptic proteins (magenta). ****P < 0.0001; Fisher’s exact test followed by Bonferroni’s correction.

**Supplementary Figure 4.**
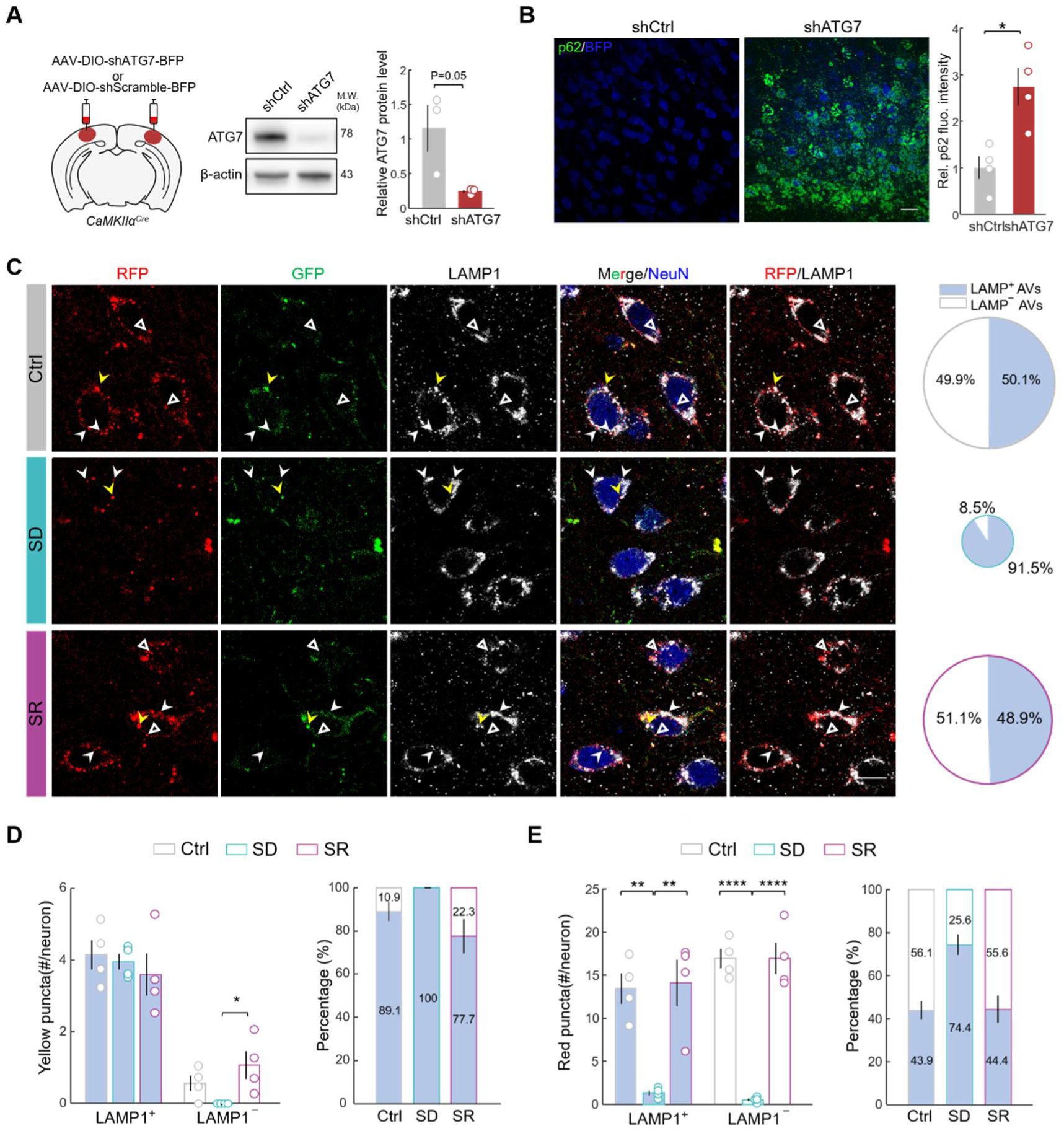
Validation of ATG7 knockdown and LAMP1 colocalization analysis in tfLC3 mice without viral manipulation. A. Immunoblot and quantification ATG7 knockdown efficiency. Each dot represents data from one mouse (N=3 mice/group). B. Representative fluorescence images and quantification of neuronal p62 levels in shCtrl and shATG7 mice. N=4 mice/group; scale bar: 20 μm. C. **Left:** Representative fluorescence images of tfLC3, LAMP1, and NeuN in the cortex of Ctrl, SD and SR mice without viral manipulation. White arrowheads: LAMP1⁺ red puncta; yellow arrowheads: LAMP1⁺ yellow puncta; triangles: LAMP1⁻ red puncta. **Right:** Pie charts showing proportions of LAMP1⁺ and LAMP1⁻ tfLC3 puncta from Ctrl (n=79 neurons, 4 mice), SD (n=138, 4 mice) and SR (n=135, 4 mice) groups. Scale bar, 10 μm. D. Abundance (left) and proportion (right) of LAMP1^+^ and LAMP1⁻ yellow puncta in cortical neurons (N=4 mice/group). Essentially all yellow puncta are LAMP1^+^. E. Similar to (D), but for red, acidified tfLC3 puncta. Data are presented as mean ± SEM. Unpaired two-tailed t-test was used in (A, B); One-way ANOVA with Bonferroni’s correction was used in (D, E). * *P < 0.05*; ** *P<0.01*; **** *P < 0.0001*.

**Supplementary Figure 5.**
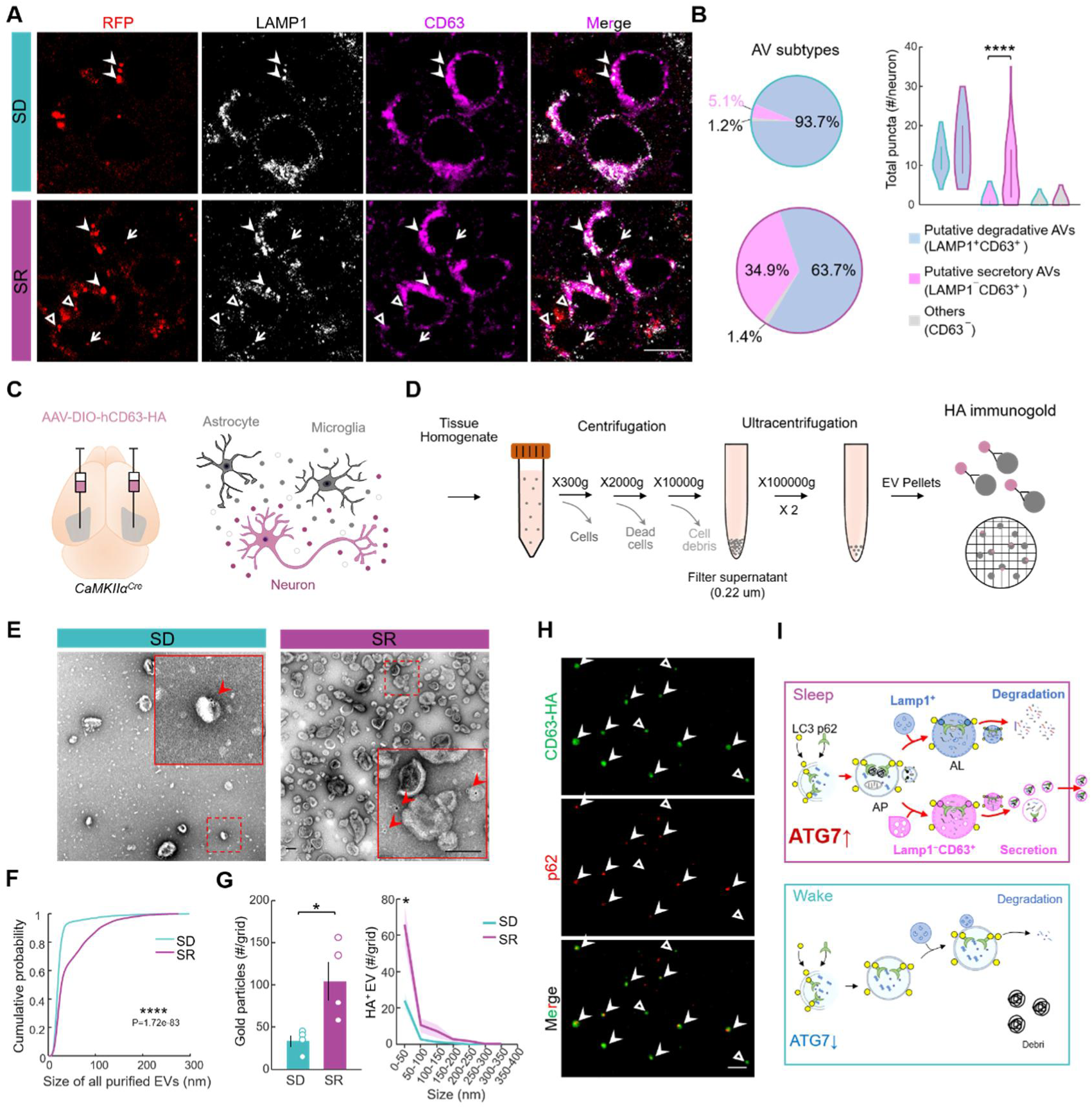
Sleep increases LAMP1⁻CD63⁺ putative secretory AVs and neuron-derived extracellular vesicles. A. Representative images showing colocalization of tfLC3 puncta with LAMP1 and CD63 (dual-markers of lysosomes and secretory vesicles). Arrowheads: representative LAMP1⁺CD63⁺ (putative degradative); triangles: LAMP1⁻CD63⁺ (putative secretory); arrows: CD63⁻ (other). Scale bar, 10 μm. B. ***Left:***Pie charts showing proportions of the three puncta types. ***Right:*** Violin plot showing the abundance of each puncta type in SD and SR (∼60 neurons from 2 mice each). Bars: IQR. C. Schematic illustrating the strategy for labeling cortical excitatory neuron-derived EVs by injecting AAV-DIO-CD63-HA into the cortex of *CaMKIIα^Cre^* mice. D. Workflow for isolation cortical EVs and detection of neuron-derived EVs by immuno-EM. E. Representative TEM images of purified cortical EVs, with HA⁺ excitatory neuron-derived EVs identified by immunogold labeling. Insets show higher-magnification views. Red arrowheads, HA^+^ gold particles. Scale bar, 100 nm. F. Cumulative size distribution of isolated EVs from SD and SR groups (∼2000 EVs pooled from 3 mice/group). G. Quantification of HA^+^ gold particle labeling in EVs (left) and their size distribution (right). Each dot represents data from one grid (N=4 grids from 3 mice/group). Error bars/shading: SEM. H. Immunostaining of HA and p62 in cortical EVs from SR mice. Arrowheads: p62⁺ HA⁺ EVs; open triangles: p62⁻ HA⁺ EVs. Note that majority of neuron-derived EVs contain p62, indicating secretion of autophagic cargo. Scale bar, 10 μm. I. Schematic summary of state-dependent autophagic pathways in cortical excitatory neurons during sleep and wake. Kruskal–Wallis test was used in B and unpaired two-tailed t-test in F. * *P* < 0.05.

**Supplementary Figure 6.**
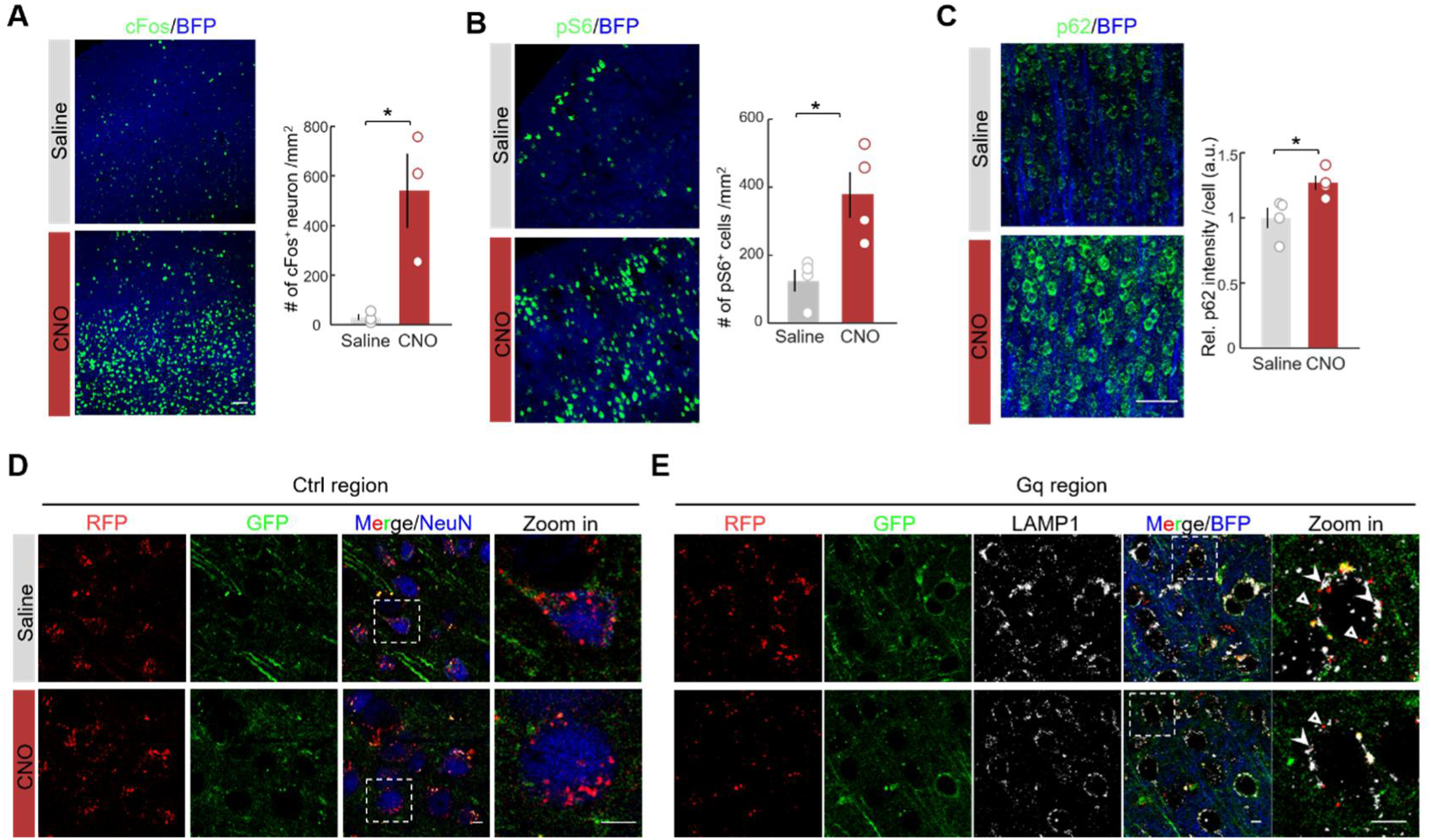
Effects of chemogenetic neuronal activation during recovery sleep on cortical autophagy. A. Representative fluorescence images and quantification of cFos expression in cortical neurons following saline or CNO treatment during SR. Each dot represents one mouse (N = 3 mice/group). Scale bar, 50 μm. B-C, Similar to A, but for phosphorylated S6 (pS6, B) and p62 (C) (N = 4 mice/group). D. Representative fluorescence images of tfLC3 puncta and NeuN in non-Gq-expressing control cortical regions. Insets show higher-magnification views. Scale bars, 5 μm. E. Representative fluorescence images showing tfLC3 puncta and LAMP1 in Gq-BFP-expressing cortical regions. Insets show higher-magnification views. Arrowheads: representative LAMP1⁺ acidified tfLC3 puncta; open triangles: LAMP1⁻ acidified tfLC3 puncta. Data are presented as mean ± SEM. Unpaired two-tailed *t* tests were used in (A-C). *\*P*<0.05.

**Supplementary Figure 7.**
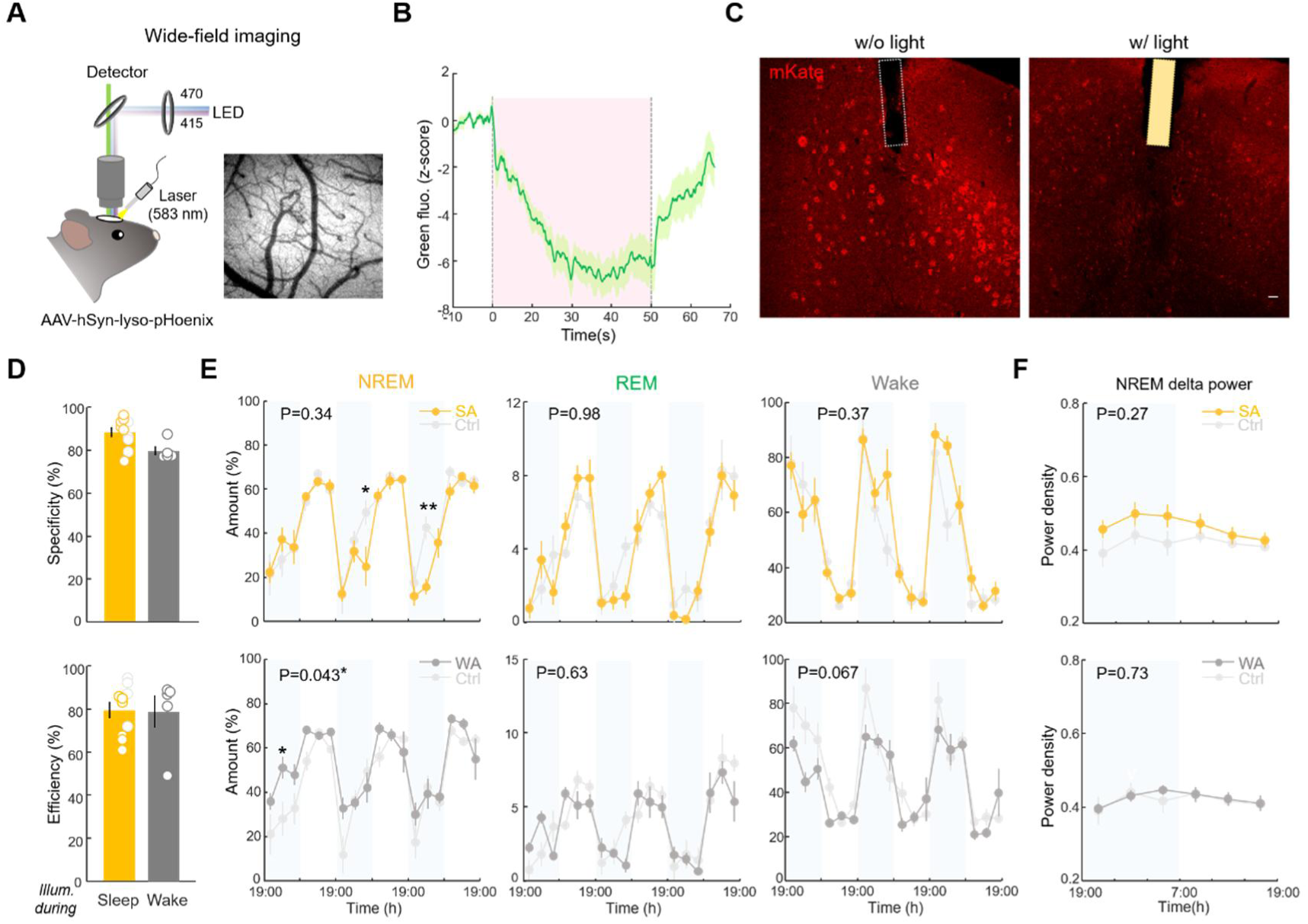
Validation of brain state-specific lysosomal acidification in vivo. A. Schematic of wide-field imaging for simultaneous Lyso-pHoenix activation and measurement of lysosomal acidification. B. Normalized pHluorin fluorescence before, during (pink shading), and after yellow-light illumination, averaged across 19 trials from 2 mice. C. Representative images of mKate fluorescence in hemispheres w/o or w/ light illumination. Reduced mKate fluorescence reflects photobleaching. Dashed outlines: optic fiber locations. Scale bar, 50 μm. D. Specificity and efficiency of closed-loop light delivery during sleep-specific acidification (SA, *N* = 6 mice), wake-specific acidification (WA, *N* = 5 mice), and sleep-specific light control (Ctrl, *N* = 4 mice). E. NREM sleep, REM sleep, and wake across three consecutive days in SA (top) and WA (bottom) mice compared with Ctrl mice. Each point represents a 4-h bin. F. Similar to (E), but for NREM delta power. Data are presented as mean ± SEM. Unpaired two-tailed *t*-tests were used in (D); two-way ANOVA followed by Bonferroni correction was used in (E, F). *P < 0.05; **P < 0.01.

**Supplementary Figure 8.**
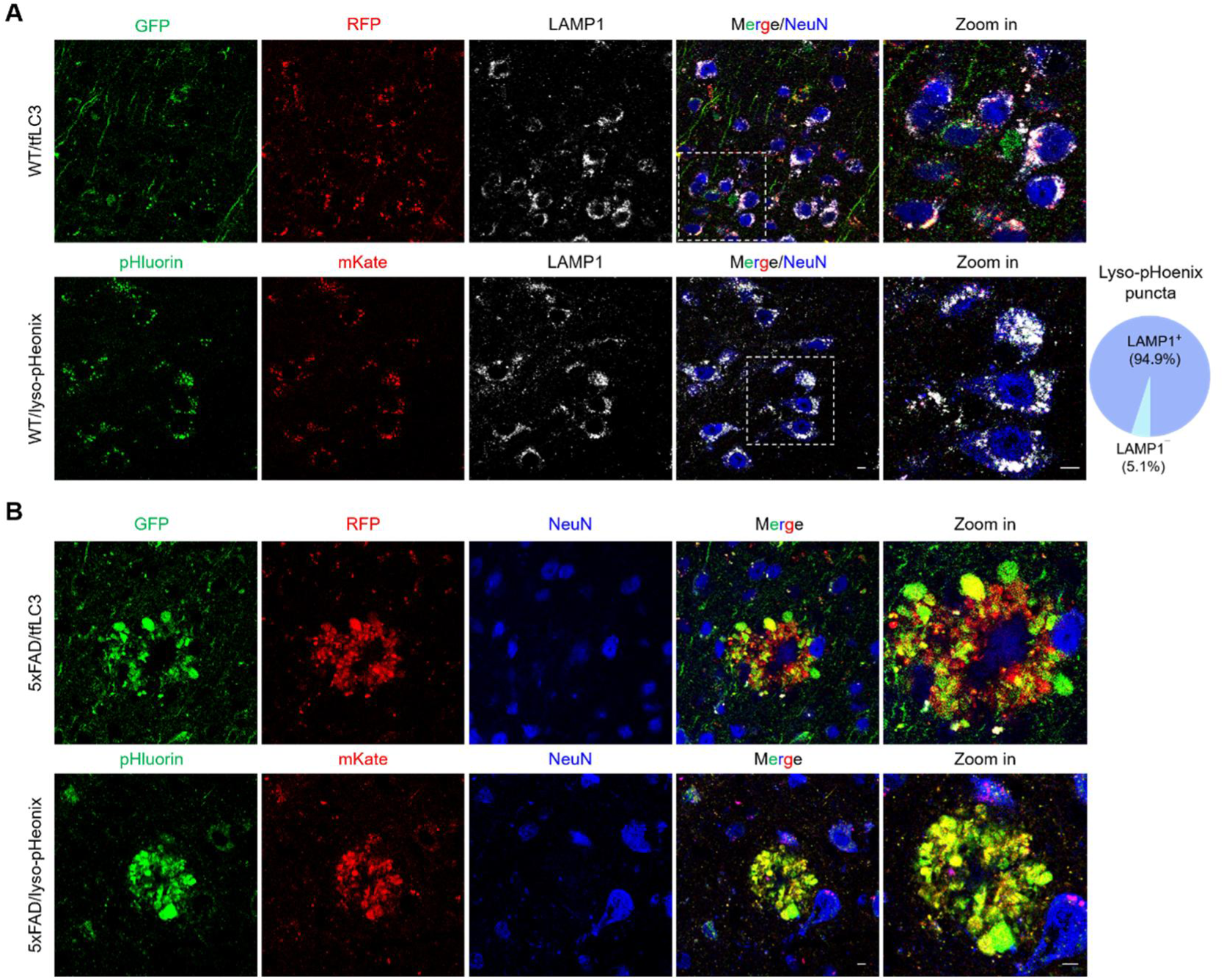
Poisonous anthos (PANTHOS) structures in 5xFAD mice visualized by tfLC3 and lyso-pHoenix. A. Representative confocal images of tfLC3 (top) or lyso-pHoenix (bottom) in wild-type mouse cortex, with LAMP1 and NeuN. Insets show enlarged views. Most Lyso-pHoenix puncta were LAMP1⁺, confirming lysosomal localization. Scale bars, 5 μm. B. Representative confocal images of tfLC3 (top) and lyso-pHoenix (bottom) in 5-month-old 5xFAD mice, showing characteristic flower-like PANTHOS structures. Scale bars, 5 μm.

**Supplementary Figure 9.**
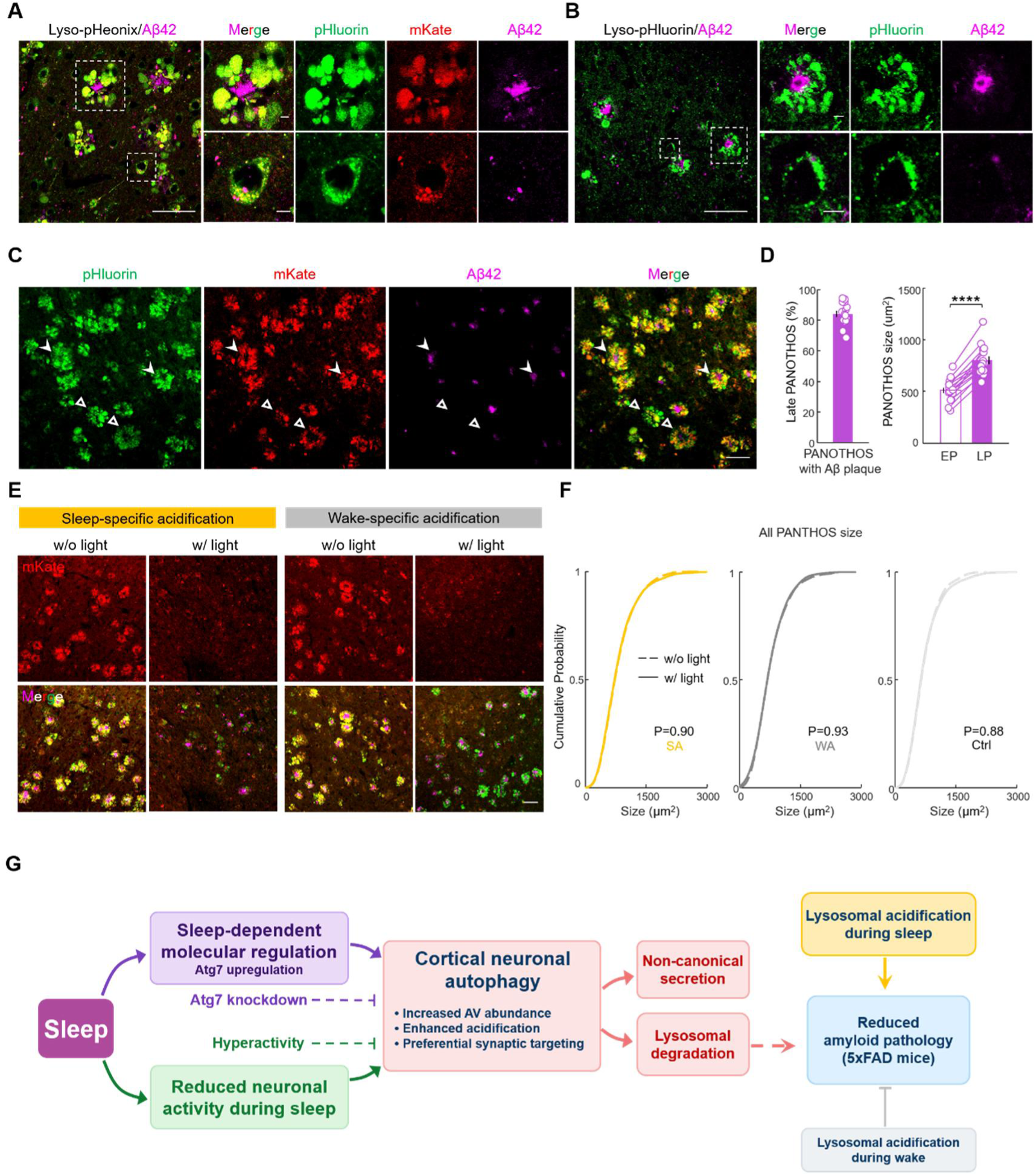
Effects of sleep- and wake-specific lysosomal acidification on PANTHOS and Aβ pathology in 5xFAD mice. A. Representative fluorescence images of Lyso-pHoenix and Aβ42 in the cortex of 5xFAD mice. Insets show a PANTHOS neuron containing Aβ42 (top) and a nearby neuron without PANTHOS (bottom). Scale bars, 50 μm (main) and 5 μm (insets). B. Similar to (A), but in Lyso-pHluorin control mice. C. Low-magnification images of Lyso-pHoenix and Aβ42 corresponding to Figure 6D. Filled arrowheads: late PANTHOS (LP); open arrowheads: early PANTHOS (EP). Scale bar, 50 μm. D. **Left,** percentage of late PANTHOS among total PANTHOS. **Right,** sizes of early and late PANTHOS. Data were pooled from non-illuminated hemispheres across SA, WA, and Ctrl mice (N = 15 mice). Each line represents one mouse. E. Representative fluorescence images of mKate, pHluorin, and Aβ42 in SA and WA mice corresponding to Figure 6C. Scale bar, 50 μm. F. Cumulative size distributions of PANTHOS in cortical hemispheres w/ (solid lines) or w/o (dashed lines) light illumination. G. Schematic summary of the proposed model. Sleep promotes cortical neuronal autophagy through sleep-dependent molecular regulation, including Atg7 upregulation, together with reduced neuronal activity. Sleep-driven autophagy preferentially targets synaptic cargo and involves lysosomal degradation together with a putative noncanonical secretory route. The efficacy of intracellular clearance depends on its alignment with sleep: in 5xFAD mice, lysosomal acidification reduced amyloid pathology during sleep but was detrimental during wakefulness. Data are presented as mean ± SEM. Paired two-tailed t-test was used in (D); Kolmogorov–Smirnov tests were used in (F). ***P < 0.001.

## Captions for Data S1

### Data S1. Proteomic profiling and annotation of LC3-immunopurified autophagic vacuoles

Included are raw proteomic measurements (n=608 proteins, from three SD and three SR replicates with paired LC3-IP and IgG-IP controls), LC3-enriched AV proteins (n=309, LC3>IgG), subcellular compartment annotations, differential cargo enrichment between SD and SR, Gene Ontology analysis, and synaptic subclassification of SR-enriched proteins.

